# Engineering a thixotropic and biochemically tunable hyaluronan and collagen bioink for biofabrication of multiple tissue construct types

**DOI:** 10.1101/2021.09.01.458584

**Authors:** Julio Aleman, Hemamylammal Sivakumar, Thomas DePalma, Yu Zhou, Andrea Mazzocchi, Richard Connor Huntwork, KyungMin Yoo, Surya Banks, Casey Clark, Alexandra Maycock, Kalan Leaks, Kevin Enck, Emmanuel C Opara, Paul Gatenholm, Mark Welker, Shay Soker, Samuel Herberg, Tracy Criswell, Aleksander Skardal

## Abstract

The field of three-dimensional (3D) bioprinting has advanced rapidly in recent years. Significant reduction in the costs associated with obtaining functional 3D bioprinting hardware platforms is both a cause and a result of these advances. As such, there are more laboratories than ever integrating bioprinting methodologies into their research. However, there is a lack of standards in the field of biofabrication governing any requirements or characteristics to support cross-compatibility with biomaterial bioinks, hardware, and different tissue types. Here we describe a modular extracellular matrix (ECM) inspired bioink comprised of collagen and hyaluronic acid base components that: 1) employ reversible internal hydrogen bonding forces to generate thixotropic materials that dynamically reduce their elastic moduli in response to increased shear stress, thus enabling increased compatibility with printing hardware; and 2) modular addons in the form of chemically-modified fibronectin and laminin that when covalently bound within the bioink support a variety of tissue types, including liver, neural, muscle, pancreatic islet, and adipose tissue. These features aim to accelerate the deployment of such bioinks for tissue engineering of functional constructs in the hands of various end users.

## Introduction

The continued advancement of three-dimensional (3D) bioprinting technologies has aided in accelerating the broader field of tissue engineering (*1, 2*). Bioprinting facilitates integration of cells, biomaterials, and other biological or synthetic compounds to fabricate customized 3D architectures that can act as surrogates of tissues or organs (*3–5*). Such bioprinted tissue constructs range in complexity from structures such as spheroids and organoids to highly complicated 3D human-scaled tissues that are being assessed for suitability for transplantation in clinical trials. While the tissue engineering field as a whole continues to encounter hurdles such as sourcing of specific cell types, mitigating risks associated with induced pluripotent stem cells, and promoting vascularization in engineered tissues of larger size, many of the technologies that have been developed within this field have evolved to the point where they have been or are being tested for use in clinical and commercial applications (*6, 7*). Bioprinting is one such technology. With the capability to drive both biofabrication of complex 3D architectures consisting of human cells and biomaterials, and to support high throughput deposition of micro-scale tissue and tumor constructs and organoids for drug screening applications, bioprinting has the potential to have a great impact outside of academia (*5, 8–13*). In recent years, an effort has been made to develop the field of regenerative medicine biomanufacturing. That is, the deployment of regenerative medicine and tissue engineering technologies and products at an industrial scale (*6, 7*). Bioprinting has been closely connected with these initiatives, yet the field of bioprinting, and biofabrication more broadly, suffers from a number of limitations, including lack of print resolution, successful printing of vasculature to support large size tissue constructs, balancing print speed, volume and resolution, and as we describe here, cross-hardware platform compatibility of bioinks and tunability of a single base bioink system to support a wide variety of tissue types (*14*). The rapid emergence of relatively affordable hardware platforms in recent years has made bioprinting more accessible than ever, which is undoubtably a positive development. However, this increased usage has produced the following problem. Put simply, there has been a tendency of researchers – new and experienced – to bioprinting, to fall back on one’s “favorite” biomaterial of the past, such as collagen type I and Matrigel, which often do not have mechanical properties conducive to the bioprinting process. We propose that more effort should be spent to develop bioinks from the ground up, based on the requirements of the dynamic processes of bioprinting and with careful consideration of the specific “needs” of the tissue being modeled. In the context of regenerative medicine biomanufacturing, we have the potential to provide user-friendly bioinks, that can be used across hardware platforms with little required customization through chemistry, modulation of viscosity, or other manipulations by the end user. A user-friendly, cross-compatible bioink should be attainable by ensuring that the bioink mechanical properties can respond dynamically to the physical forces imbued upon it by the printer, and be tuned biologically without affecting printability (*14*).

Since bioprinting emerged in the 2000s, the realization of the technology has been plagued by the concept of the “printability window”. In the context of additive manufacturing, the timeframe to crosslink a bioink between extrusion and deposition is defined by the initial state of the carrier polymer. To form a stable 3D structure, a crosslinking reaction must occur. The posed problem is: how does one transition a semisolid cell suspension of a hydrogel precursor, move it through a bioprinter printhead, and have it hold its predetermined 3D geometry post-deposition all while not negatively affecting cell viability? Generally, printing a material too early results in a puddle of uncrosslinked polymer and cells. On the other hand, if the material is printed too late, the bioink may crosslink to the extent where either it extrudes, but fractures as it passes through a printhead, or it completely occludes the printer. Our early efforts focused on solving this problem in the context of hyaluronic acid (HA) and gelatin hydrogels. We were able to extend the bioprinting window by using stepwise UV crosslinking, thus creating partially crosslinked networks of methacrylated HA and gelatin that were extrudable but held their shape until they could be fused post printing by a second UV light exposure (*15*). In another strategy, we employed slower crosslinking thiolated HA and gelatin materials combined with gold nanoparticles as crosslinkers, thus harnessing gold-thiol interactions to generate an 8-16 hour biofabrication window (*16*). Unfortunately, the use of nanoparticles entails a potential future clinical hurdle, limiting the utility of this method for translational applications. Subsequently, we employed bioinks based on discrete multiple chemical reactions to again create steady state biofabrication windows and leverage secondary reactions for solidifying the printed construct or further increasing its elastic modulus to match specific tissue elastic moduli (*17–19*).

Many components of the extracellular matrix (ECM) are crucial to biological function, and as such should not be ignored as bioprinting, biofabrication, and tissue engineering enter a new period in which they are more acutely assessed for translational applications and eventually biomanufacturing (*6, 7*). Materials such as Matrigel and decellularized ECM hydrogels are likely non-starters due to their sourcing and undefined composition (*20*). Rather, a bioink system composed of defined natural materials that are modified synthetically in which the addition of key ECM components can be tuned based on the desired tissue type while printability is controlled through chemistry native to that bioink, as opposed to end user customization, has the potential to facilitate increased adoption of the technology by those end users and spur an effort at standardizing bioink specifications in bioprinting.

In this work, we address these shortcomings by implementing a “tool kit” that supports a thixotropic ECM biomimetic HA and collagen bioink that is easily printable. Chemically modified ECM proteins (fibronectin [FN] and laminin [LMN]) are covalently bound to the hydrogel network to support the creation of a variety of types of tissue constructs. We employ catecholamine groups to induce hydrogen bonding within the bioinks, which is reversible and can be tuned to create a thixotropic material. In response to the shear stresses during bioprinting, this thixotropic material dynamically undergoes a decrease in elastic modulus allowing printing to occur. The elastic modulus rebounds after passing through the shear stress inducing printhead, allowing the bioink to hold its shape. From a biochemical standpoint, modulation of HA, collagen, FN, and LMN provide a toolbox of ECM signals that, while simplistic compared to Matrigel and whole tissue decellularized ECM, is well-defined and still supports the growth of functional tissue constructs, as we demonstrate.

Taken together, the studies herein provide a major advancement towards standardization of bioprinting and bioink technology that more effectively addresses the biofabrication window problem compared to previous efforts. At the same time, we demonstrate that a relatively small set of key ECM components (HA, collagen, FN, and LMN) can be combined to support a wide variety of cell types and tissues, including liver, neural, pancreas, adipose, and skeletal muscle. Leveraging this and other comparable technologies may allow our field to come closer to realizing the long-held potential of tissue engineering and 3D bioprinting.

## Results

### Bioink Strategy

HA and gelatin-based hydrogels have served our laboratory’s interests for many years. However, in studies over the past several years, it became clear that the presence of fibrillar collagen can be a crucial component to mimicking *in vivo* biology *in vitro* (*21, 22*). In both healthy and malignant tissues, collagen alignment can play a significant role in the induction of pathways that control cell fate (*23–25*). Based on these observations, we recently described an HA-collagen bioink that while effective in some applications, suffered from the “biofabrication or printability window” (i.e., the timeframe to crosslink the bioink between extrusion and deposition) described above. Thus, we sought to develop a strategy (***Fig. 1a-c***) that could be applied to our HA-collagen bioink – or any bioink, including our former HA-gelatin varieties) to improve printability and remove as much user error from the process as possible. This platform technology is based on methacrylated collagen (Coll) and thiolated-HA (***Fig. 1a***), which will crosslink directly or can be modified with alternative crosslinking (hydrogen-bonding [H-bond]) capabilities for reversible crosslinking/thixotropic properties to facilitate straightforward extrusion bioprinting. Modification of cell adherent proteins or peptides with functional groups supports cell adhesion and can be varied based on specific tissue characteristics (***Fig. 1b***). Additionally, heparin/heparan sulfate pendant chains allow for growth factor immobilization, if desired. Lastly, while not a primary focus herein, the balance of thiol, methacrylate, and catecholamine groups can be tuned to drive overall bulk elastic modulus changes to match desired tissue mechanical properties (***Fig. 1c***).

**Figure 1.**
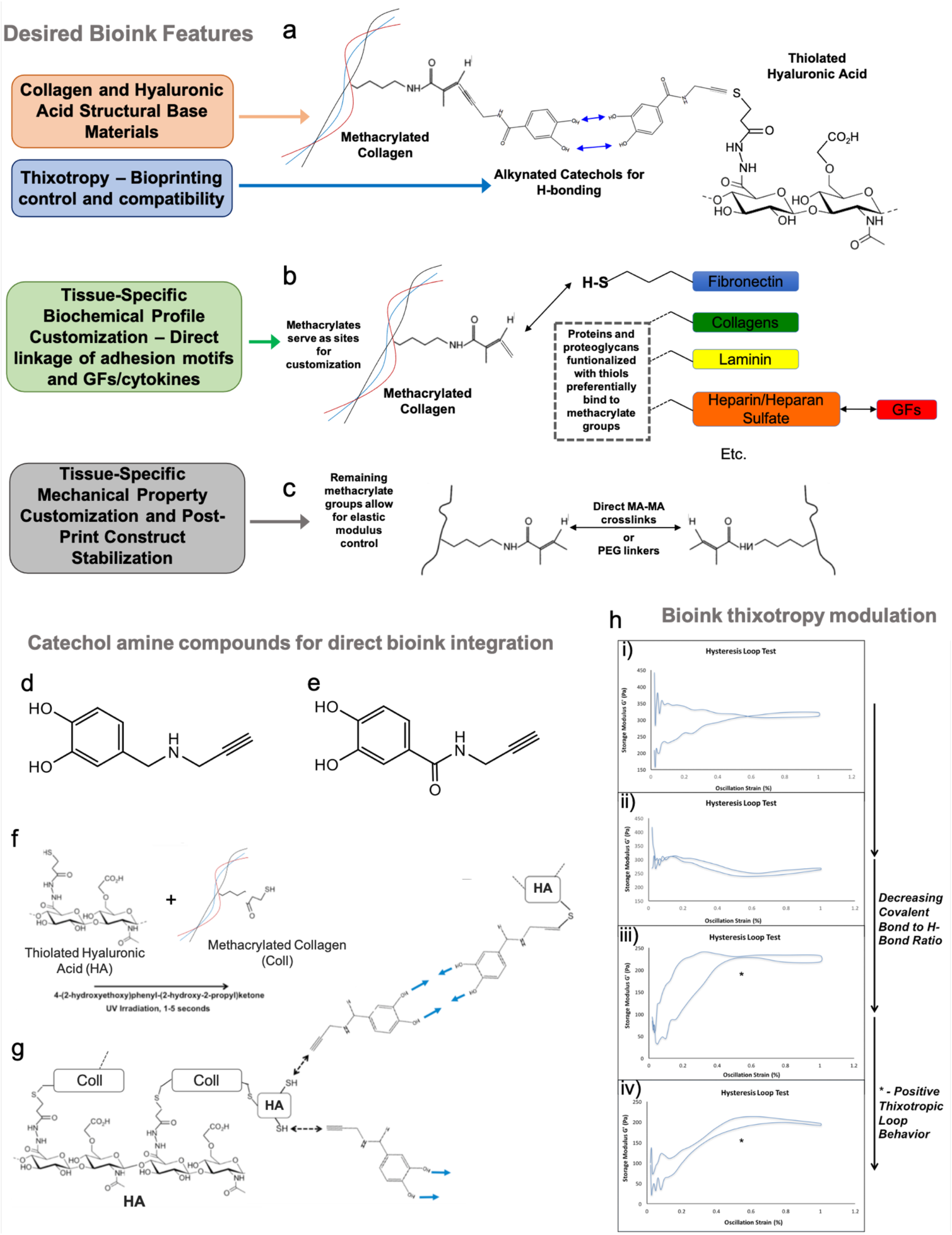
Generating a highly functional and usable ECM bionk requires control over both biochemical profiles and mechanical properties. a-c) The desired features of a versatile bioink include: a) Natural ECM-derived base components, realized through the use of methacrylated collagen and thiolated hyaluronic acid. In addition, modulation of crosslinking methods can imbue thixotropic properties to the bioink through the use of reversible hydrogen bonding. b) Customizable biochemical profiles, attained through the use of thiolated adhesion proteins and optional growth factor immobilization through heparin pendant chains on the hyaluronic acid component. c) Tissue-specific mechanical property customization through a second, optional crosslinking step. d-h) Incorporation of alkynated catecholamines within the bioink enables thixotropic mechanical properties through hydrogen bonding. d-e) Chemical structures of the catecholamine addons. The compound shown in panel d) was deployed in subsequent studies. f-g) incorporation into a hyaluronic acid hydrogel system via a thiol-alkyne reaction. h) By modulating the amount of catecholamine, and thereby modulating the ratio of covalent bonds to H-bonds, one can tune the bioink from h(i-ii)) non-thixotropic materials to h(iii-iv) thixotropic materials that extrude smoothly.

### Design and testing of an ECM-based catecholamine-enabled thixotropic bioink

Thixotropic hydrogels have been developed previously, but it has been difficult to imbue within natural ECM-derived biomaterials. In thixotropic materials, the elastic modulus decreases in response to external force but is restored once that force is removed. Covalently conjugating functionalized catecholamine small molecules (***Fig. 1d-e***) (*26*) to methacrylated-Coll and thiolated-HA precursors produces a thixotropic material because reversable H-bonds are formed between the alcohol groups of adjacent catechols (***Fig. 1f-g***).

Through analysis of the subsequent hydrogels, we found that we could modulate thixotropic properties by decreasing or increasing the ratios of covalent bonds (methacrylate-methacrylate or methacrylate-S vs. H-bonds), inducing clear differences in thixotropy hysteresis loop rheological tests (***Fig. 1h(i-iv)***). Low H-bond bioinks exhibited non-thixotropic behavior (***Fig. 1h(i)***), whereas hysteresis tests showed the production of principally thixotropic hysteresis loops with increasing H-bond activity (***Fig. 1h(iii-iv)***). This suggests that these bioinks only need one “recovery” or “thixotropic cycle” of elastic modulus to be a useful bioink.

To assess practical printability, the HA-Coll bioink was tested using different size nozzles and speeds, thus optimizing print speed and resolution. ***Fig. S1*** describes the effects of speed input on print time and resolution for 23-gauge and 30-gauge printhead nozzle tips while printing a simple 3-segment hydrogel construct. For both tip sizes there was an expected decrease in print time as print speed increased (***Fig. S1a-b***). However, print resolution and aesthetics varied across print speeds. As we observed visually, thickness of hydrogel filaments decreased with increasing print speed (***Fig. S1e***). For 23-gauge tips, consistency of the filaments deteriorated above 20 mm/s. For 30-gauge tips, consistency was preserved more effectively. Consistency in resolution was quantified, after which the standard deviation of filament width was determined. Again, 23-gauge tips showed more variation, while for 30-gauge tips, several print speeds (24 mm/s, 32 mm/s, and 36 mm/s) minimized standard deviation, while also minimizing print time (***Fig. S1c-d***). Of these, 24 mm/s striked the best balance between print time, resolution, and visual consistency of the printed filaments, and as such, was employed in subsequent bioprinting protocols.

### Integration of covalently bound fibronectin and laminin can drive cellular phenotype

Biomaterial approaches to tissue engineering have largely skewed towards using overly simplistic matrices alone (gelatin, alginate, HA, polyethylene glycol diacrylate (PEGDA), etc…) or matrices modified with adhesion peptides. Native tissue ECM is considerably more complex and varies between tissue types. Therefore, one-size-fits-all solutions like Matrigel are not ideal, nor is Matrigel translatable for clinical applications. Decellularized ECM materials suffer from similar problems, as the presence of various growth factors, cytokines, and other components vary from batch to batch and are difficult to control. Instead, we believe that a ground up approach to ECM engineering is the solution. We selected the key ECM proteins FN and LMN and functionalized them with thiol groups so that they could be covalently crosslinked into our bioinks (***Fig. 2a-b, Fig. S2a***). First, we assessed the effect of covalently bound FN on cell adherence. To do this, the HCT-116 colorectal cancer cell line was seeded atop preformed HA+PEGDA, HA+FN+PEGDA, or HA+Gelatin+PEGDA (***Fig. S2b***). On HA+PEGDA, a bioink lacking cell adhesion peptide sequences, cells aggregated with one another, but did not adhere to the hydrogel, and were easily washed away during routine phalloidin staining. By contrast, on both HA+FN+PEGDA and HA+Gelatin+PEGDA bioinks, which provide cell adhesion motifs by either the covalently bound FN or gelatin, cells robustly attached to the hydrogels. In a second validation study designed to verify the need for covalent binding of FN to the matrix, human umbilical vein endothelial cells (HUVECs) were plated on HA+PEGDA, HA+PEGDA with FN added in the media, HA+PEGDA with un-thiolated FN added in the hydrogel, or HA+FN+PEGDA (***Fig. S2c***). Only in the HA+FN+PEGDA, in which FN is stably coupled to the matrix, did the cells adhere to the hydrogel. In the other 3 conditions, the HUVECs did not attach to the hydrogels and instead aggregated with one another.

**Figure 2.**
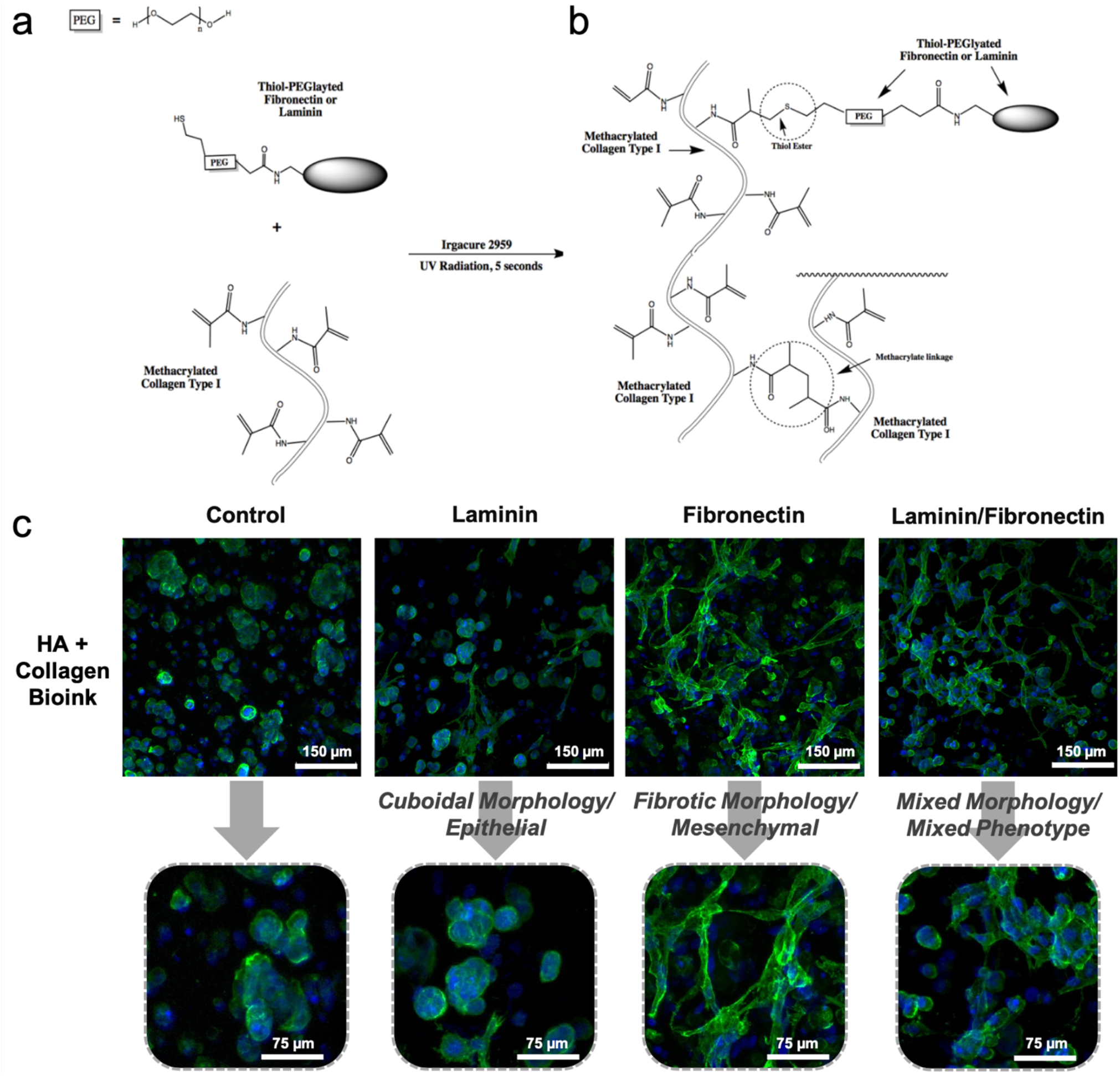
Functionalization of adhesion proteins - laminin and fibronectin - and incorporation into 3D tissue constructs drives cell phenotype. a-b) Integration of thiolated adhesion proteins into HA-Coll hydrogels via thiol-methacrylate linkages. c) LX2 stellate cells are shown via macro-confocal microscopy following staining with phalloidin (green) and DAPI (blue). The addition of thiolated laminin conserves an epithelial phenotype. The addition of thiolated fibronectin drives many cells toward a mesenchymal phenotype. A 1:1 ratio of laminin and fibronectin results in a mixed phenotype. Scale bars – 75 or 150 μm.

Additional viability and proliferation assays were used to show that the addition of the chemically modified adhesion protein (in this case FN) did not cause any cell toxicity due to potential unreacted reagents or byproducts. As shown in ***Fig. S3a-f***, using HepG2 liver hepatoma cells, U87 MG glioma cells and HCT-116 cells, the addition of modified FN did not negatively impact viability or proliferation of these cell lines, and in fact appeared to be a positive influence on both metrics. It should be noted that these cell lines are robust; thus, viability of additional cell types was verified subsequently throughout this study.

Lastly, the influence of ECM component modulation on general cell phenotype was determined by assessing the morphology of LX2 hepatic stellate cells within bioink formulations. Confocal imaging of the stellate cells showed a correlation between morphology and adhesion protein content within hydrogels (***Fig. 2c***). When compared to the HA+Coll base bioink, the addition of LMN resulted in minor changes to cell morphology, with cells largely remaining more epithelial-like and cuboidal. Conversely, adding FN alone drove cells to take on a more pronounced mesenchymal and potentially fibrotic morphology based on the elongated and branched morphology of the stellae cells. The addition of both LMN and FN caused cells to display a mixture of these morphologies. While these output metrics are only qualitative, they suggested the potency of using only a handful of physical, ECM-based signals to support different distinct cell and perhaps tissue phenotypes, which we further explored in the remainder of this work.

### Hydrogel characterization following covalent incorporation of adhesion proteins

First, the HA-Coll and HA-Coll-FN hydrogels were characterized by rheological testing, using a commercially available HA-Gel hydrogel (i.e., Hystem) as a comparison. Strain sweep tests (0 to 1000% strain) showed that the addition of FN to the HA-Coll hydrogels did not significantly change the mechanical properties of the material. Both HA-Coll and HA-Coll-FN had elastic moduli in the range of 500 to 1000 Pa (***Fig. S4a and S4b***). Moreover, both formulations also shared a similar failure point at approximately 300% strain. In comparison, the control HA-Gel hydrogel had an elastic modulus in the 100 to 250 Pa range (***Fig. S4c***), consistent with our previous studies (*16, 18, 27, 28*). While not addressed explicitly here, elsewhere we have shown the elastic modulus of HA-Coll hydrogels can be further tuned by modulating protein/polymer concentration and crosslinking methods (*22*).

Next, porosity of these 3 hydrogel formulations was determined by analyzing scanning electron microscopy images (***Fig. S4d-f***). Visually, as well as quantitatively (***Fig. S4g***), the addition of FN did not appear to change the average pore size (about 60 μm in diameter). In comparison, the HA-Gel hydrogels had significantly larger pores (approximately 130 μm in diameter). This is expected as the HA-Gel has a lower elastic modulus, which typically corresponds to a looser crosslinking density, and thus increased porosity.

Additionally, swelling behavior was assessed to determine whether or not the hydrogels would maintain a consistent volume and fluid content over time. ***Fig. S4h*** shows raw weight data, including initial weight and weight after 24 hours incubation in DI water. All hydrogel formulations tested showed relatively minimal levels of swelling. HA-Coll hydrogels swelled 10.3%, gaining 9 mg in water weight on average. Conversely, HA-Coll-FN hydrogels contracted slightly, by 7%, suggesting that the inclusion of FN may result in some internal interactions, either through hydrophobic interactions or a low number of instances in which the thiolated FN proteins serve as additional crosslinkers between HA and collagen components. In comparison, the HA-Gel swelled by 13%, gaining 11 mg in water weight on average. These swelling and contraction values fall well within ranges described in previous studies that evaluated both commercially available, widely accepted, or experimental hydrogel biomaterials (*29*).

### Liver construct engineering

To date, primary human hepatocyte cultures continue to be the gold standard for liver-specific toxicology screening *in vitro*. We previously showed the potency of using decellularized and dissolved liver tissue ECM as a supplement in hydrogels to successfully support hepatocytes (*30*). While this approach did not use tumor-derived ECM as in Matrigel, it still suffers from the black box nature of tissue ECM products (i.e., uncharacterized cytokines, growth factors, and other components) as well as batch to batch variability. Using our defined bioink system, we explored its suitability for 3D primary hepatocyte cultures for toxicology testing. The simple inclusion of FN in the HA-Collagen bioink resulted in much improved long-term viability of hepatocytes in 3D culture via LIVE/DEAD staining (***Fig. 3a***). Likewise, we observed significantly increased albumin and urea secretion for over 2 weeks by hepatocytes in FN-supplemented bioinks (***Fig. 3b-c***).

**Figure 3.**
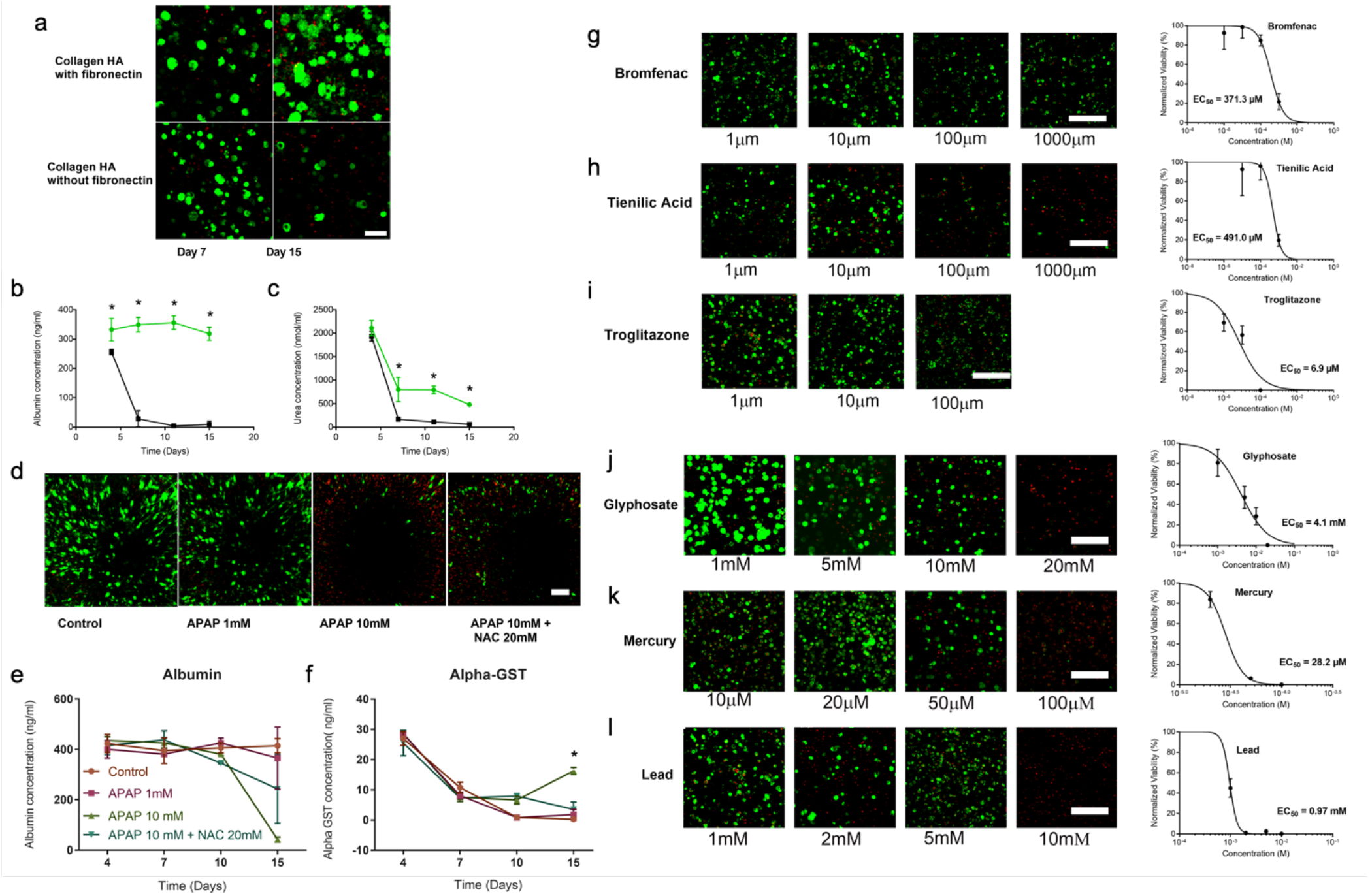
Bioink incorporation of fibronectin increases basic primary hepatocyte-based liver construct function and facilitates functional drug screening assays. a-c) Viability and albumin and urea output. a) Representative live/dead images (live = green cells; red = dead cell nuclei) of liver organoids in the HA-Coll and HA-Coll-FN hydrogels on days 7 and 15. Scale bar – 100 μm. Quantification of b) albumin (ng/ml) and c) urea (nmol/ml) from liver organoids in collagen gel formulation with and without fibronectin at days 4, 7, 11, and 15 (n = 3 per time point). * represents statistical significance corresponding to P value less than 0.05. d-f) APAP toxicity and NAC rescue with liver organoids. d) Representative live/dead images (live = green cells; red = dead cells) Scale bar – 200 μm. e-f) Albumin and alpha-GST secretion of the liver organoids on day 4,7,10 and 15, as measured by ELISA. Statistical significance: * p < 0.05 between APAP 10 mM condition and all other conditions. g-l) Toxicity screens using FDA recalled drugs and environmental toxins. Representative live/dead images (live = green cells; red = dead cells) of liver organoids in HA-Coll-FN hydrogels exposed for 48 hours to the FDA recalled drugs g) bromfenac sodium at concentrations of 1 μM, 10 μm, 100 μM, and 1000 μM, h) tienilic acid at concentrations of 1 μM, 10 μm, 100 μM, and 1000 μM, and i) troglitazone at concentrations of 1 μm, 10 μM, and 100 μM. Scale bars – 200 μm. Half-maximal effective concentrations (EC_50_) of g) bromfenac sodium, h) tienilic acid, and i) troglitazone (n = 6 per concentration) as determined from ATP activity curves normalized to control liver organoids without drug. Analogous data for environmental toxins j) glyphosate at concentrations of 1 mM, 2 mM, 5 mM, and 20mM, k) mercury chloride at concentrations of 10 μM, 20 μM, 50 μM, and 100 μM, and l) lead chloride at concentrations of 1 mM, 2 mM, 5 mM and 10 mM. Scale bars – 100 μM. Half-maximal effective concentrations (EC_50_) of j) glyphosate, k) mercury chloride, and l) lead chloride (n = 6 per concentration) as determined from ATP activity curves normalized to control liver organoids without drug.

Next, to better understand hepatocyte function in our bioink, we performed a toxicity experiment that has been used in previous studies. We induced liver toxicity in our 3D liver constructs with acetaminophen (APAP) and provided N-acetyl-L-cysteine (NAC) as the clinically approved countermeasure, as we have described previously in studies with primary liver spheroids (*11*). LIVE/DEAD staining showed increased hepatocyte death with increase APAP concentration (1 mM to 10 mM) while treatment with 20 mM NAC appeared to impede or slow the toxic effects of APAP (***Fig. 3d***). While this reduction in toxicity was slight, it was supported by quantification of albumin (biomarker of liver function, which was increased with NAC treatment) and alpha-glutathione-S-transferase (biomarker of liver cell death, partially mitigated with NAC treatment) (***Fig. 3e-f***).

The drugs bromfenac sodium, troglitazone, and tienilic acid, recalled by the FDA for hepatotoxicity, were investigated (*31, 32*). LIVE/DEAD analysis of HA-Coll-FN liver constructs in presence of increasing concentrations of bromfenac sodium demonstrated increased cell death in a dose dependent manner (***Fig. 3g***), with near to complete cell death observed at 1000 μM. The half-maximal effective concentration (EC_50_) of bromfenac sodium was 371.3 μM (***Fig. 3g***). Comparable LIVE/DEAD staining patterns were noted with troglitazone and tienilic acid (**Fig. 3h, i**), and analysis of cell death revealed an EC_50_ of 6.9 μM for troglitazone and 491.0 μM for tienilic acid, respectively. This data suggests that bioengineered liver constructs exhibit dose-dependent sensitivity to drugs recalled by the FDA, thereby validating our model.

Next, the response of liver constructs exposed to various environmental toxins with known hepatotoxicity for 48 hours was investigated. LIVE/DEAD analysis of HA-Coll-FN liver constructs in the presence of increasing concentrations of glyphosate demonstrated increased cell death in a dose dependent manner (***Fig. 3j***), with near to complete cell death observed at 20 mM. The EC_50_ of glyphosate was 4.1 mM (***Fig. 3j***). Comparable LIVE/DEAD staining patterns were noted with mercury chloride and lead chloride (***Fig. 3k, l***), and analysis of cell death revealed an EC_50_ of 28.2 μM for mercury chloride and 0.97 mM for lead chloride, respectively (***Fig. 3k, l***). This data suggests that bioengineered liver constructs exhibit dose-dependent sensitivity to environmental toxins with known hepatotoxic effects.

### Hydrogel bioinks support engineered neural constructs and blood brain barrier models

We employed a similar approach to assessing the utility of our bioink platform in the context of the central nervous system. Specifically, we used primary human mesenchymal stem cells to derive astrocytes, as well as engineer a simplistic blood brain barrier model. Of note, for the neural tissue models, we specifically avoided using type I collagen, as it is not widely present in the brain. Instead, together with thiolated HA, we used the thiolated gelatin and PEGDA platform employed in a number of past studies, which is also compatible with 3D bioprinting, (*28, 33–37*) modulated with thiolated LMN and/or FN (***Fig. S5a***). Initial assessment of the neural environment was performed using co-cultures of astrocytes, human brain microvascular pericytes, and human brain endothelial cells in bioink constructs. All hydrogel formulations successfully supported viable cultures at 7 days (***Fig. S5b***). Neurons have always been challenging to maintain with an appropriate phenotype in most 3D *in vitro* environments (*38*). Here, we also used morphology as an assessment tool. We screened a matrix of bioink formulations (control, +FN, +LMN, and +FN+LMN) versus several stiffness ranges relevant to neural biology: 200-400 Pa (linear crosslinker, low end of brain stiffness), 1000 Pa (50:50 linear:4-arm crosslinker, high end of brain stiffness) and 2000 Pa (4-arm crosslinker, greater than brain stiffness). We used LIVE/DEAD staining to qualitatively evaluate both viability and morphology (***Fig. 4a-c***). Several trends emerged. First, LMN drove enlarged cell bodies and a morphology not indicative of neurons. This was also observed, but to a lesser extent in LMN+FN cultures. Stiffness also played a role as increased stiffness above that of most brain tissue, measured by rheology, resulted in elevated cell death and fewer axonal projections. A FN-modified HA-gelatin bioink at a stiffness of approximately 200-400 Pa generated the most evident extended cell morphology, suggestive of an appropriate microenvironment to start from for future neuronal cultures. In addition, astrocytes also exhibited greater cell spreading in FN-modified bioinks versus LMN-modified bioinks (***Fig. 4d-e)***.

**Figure 4.**
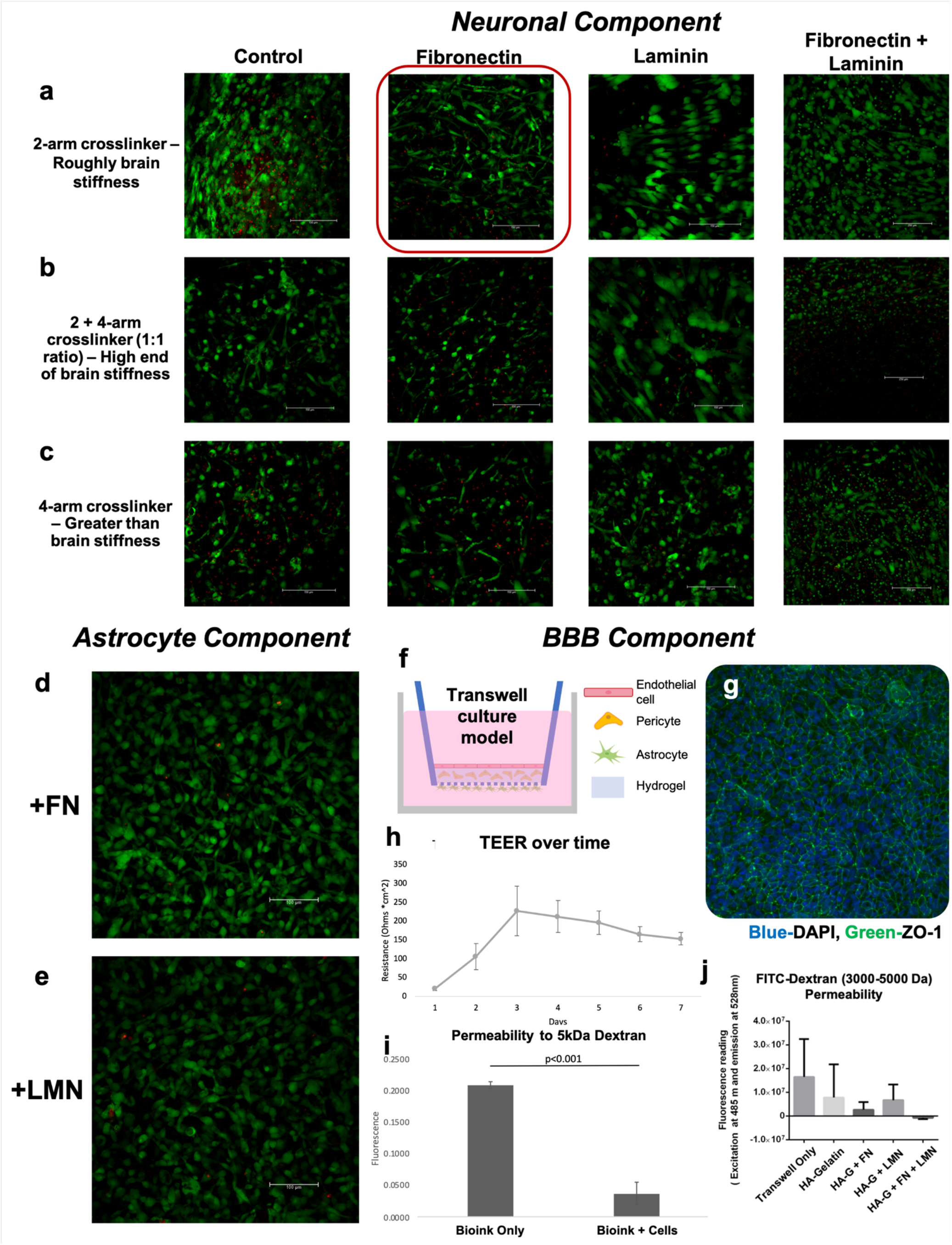
ECM component modulation drives neural cell morphology and blood brain barrier model permeability. a-c) MSC-derived neuron-like cells encapsulated in HA-gelatin (control), +FN, +LMN, or +FN+LMN bioinks, which were modulated to a) brain stiffness, b) the high end of brain stiffness, or c) greater than brain stiffness through the use of PEG-based crosslinkers of varying geometries. LIVE/DEAD staining and macro-confocal imaging was used to visual cellular morphology, as well as viability. Scale bars – 150 μm. Red box indicates the condition with the highest degree of cellular extension. d-e) Astrocytes maintained within bioinks supplemented with fibronectin or laminin, also subjected to LIVE/DEAD staining and macro-confocal imaging for visualization. Scale bars – 100 μm. Green – Calcein AM-stained viable cells; Red – Ethidium homodimer-1-stained dead cell nuclei. f-k) Blood brain barrier (BBB) model evaluation. f) Schematic of the layered Transwell BBB model, containing pericytes embedded within the hydrogel bioink over the porous membrane. Astrocytes are seeded on the other side of the membrane, while endothelial cells populate the surface of the bioink inside the insert. Within several days g) immunofluorescent staining indicates a robust endothelium (green – ZO-1; blue – DAPI) and h) trans-endothelial electrical resistance (TEER) sensing indicates increasing TEER as confluency increases, after which TEER levels off. i-j) BBB permeability evaluation. i) Permeability of a base hydrogel only control and a cellularized base hydrogel to 5 kDa FITC-dextran. Permeability in cellularized bioink constructs is significantly decreased (p < 0.001). j) Permeability of ECM bioinks to 3000-5000 kDa FITC-dextran as a result of ECM component modulation.

The blood brain barrier (BBB) prevents toxins from entering the brain. It can also serve as a barrier to otherwise effective drug compounds. Understanding and correctly modeling this critically important feature of the central nervous system (CNS) will be crucial for future drug development in the context of CNS-based disease. We used a simple Transwell BBB model to evaluate bioink composition (***Fig. 4f***). Our platform achieved a robust and confluent BBB endothelium (***Fig. 4g***), maintenance of trans-endothelial electrical resistance (***Fig. 4h***), and slowed dextran transport (***Fig. 4i***), which was further modulated by ECM composition supporting the BBB model (***Fig. 4j***). Interestingly, the most effective BBB ECM formulation included LMN and FN, which unlike the neuronal cultures, appeared to benefit from the addition of FN and not LMN. Using a FITC-dextran mass transport assay, we observed that the BBB construct formed using the HA+gelatin bioink with added FN and LMN prevented transport of the dextran (MW 3000-5000 Da) through the barriers and into the wells (***Fig. 4j***). In comparison, all other BBB constructs (HA+gelatin and HA+gelatin with FN or LMN only) allowed dextran to pass through the barriers.

### Skeletal muscle construct engineering

For creating skeletal muscle constructs, previous research had shown success in achieving proper cell phenotype without needing to manipulate specific ECM adhesion proteins. However, it has been challenging to engineer a system that promotes myocyte alignment and fusion within a 3D volume (*39*). For example, in preliminary studies using C2C12 murine myoblasts, when formed into cell spheroids, there is little cell fusion and no apparent alignment (***Fig. S6a-b***). Since it is known that physical signals such as fiber direction and topography can drive myocyte alignment and fusion, we wanted to explore the concept of using the physical bioprinted bioink filaments themselves to drive this behavior (***Fig. S6c***). Two printhead nozzle sizes were used – a larger 1 mm diameter nozzle and a smaller 0.3 mm nozzle. Interestingly, some cell fusion was observed in the larger diameter filaments after 9 days of culture after differentiation media was added, while no fusion was observed in the smaller diameter filaments (***Fig. S6d***). However, no cell alignment was observed in any of the conditions.

Given the lack of cell alignment and inconsistent cell fusion, we next employed an alternative strategy to reach the desired morphology and phenotype through manipulation of hydrogel topography. Using a polydimethylsiloxane (PDMS) stamp with 100 μm parallel ridges set 200 μm apart (***Fig. S7a(i-iii)***), parallel grooves with a thin layer of bioink were patterned by stamp placement over the bioink layers following a crosslinking step, after which C2C12 cells were incorporated (***Fig. S7a(iv-vii)***). After several days, alignment and fusion were observed, and this trend was pronounced with the addition of differentiation media (***Fig. S7b(i-iv)***). Importantly, in comparison with unpatterned bioinks or 2D culture on tissue culture plastic, we observed a distinct difference in alignment, with significantly increased cell alignment on the micropatterned bioinks (***Fig. S7c(i-iii)***). Cell fusion was observed in both conditions. When these constructs were stained for myosin heavy chain (MHC) and DAPI, we observed the same clear trend in cell alignment, as well as cell fusion in micropatterned constructs (***Fig. S7d(i)***), and what appeared to be less consistent cell elongation and fusion in the unpatterned constructs (***Fig. S7d(ii)***).

To demonstrate translational impact, this micropatterning approach was then deployed using primary human skeletal muscle progenitor cells. Within 5 days of culture in differentiation media we observed robust cell alignment and fusion in the 3D micropatterned condition, and this phenotype continued throughout day 14 (***Fig. 5a***). In comparison, the unpatterned bioink supported some fusion and regional, yet inconsistent, alignment. First, MHC expression was assessed in the 3D versus 2D conditions, under LG and HG media conditions. Fluorescent imaging showed that regardless of glucose level, the 3D micropatterned bioink yielded consistent aligned and fused cellular constructs (***Fig. 5b***). Unpatterned bioinks yielded fused, but unaligned cell organizations. Cell culture media aliquots were collected on day 2 and day 4 following LG versus HG challenge, and were assessed for concentrations of secreted interleukin (IL)-6 and GDF-8 (myostatin) (***Fig. 5c-d***). Both proteins have been shown to play roles in the crosstalk between muscle, pancreas, and fat during dramatic changes in serum glucose levels, such as those experienced by some diabetic subjects (*40–43*). Interestingly, we observed a more dynamic IL-6 response in 3D micropatterned constructs, where there was a significant difference between LG and HG-induced levels of IL-6, initially increased in HG conditions on day 2, after which LG conditioned constructs increased IL-6 concentrations significantly by day 4. In contrast, unpatterned constructs showed minimal responses to changes in glucose levels. GDF-8, on the other hand, was maintained at a consistent level in 3D micropatterned constructs regardless of glucose level, but decreased in unpatterned HG conditions, but only on day 2.

**Figure 5.**
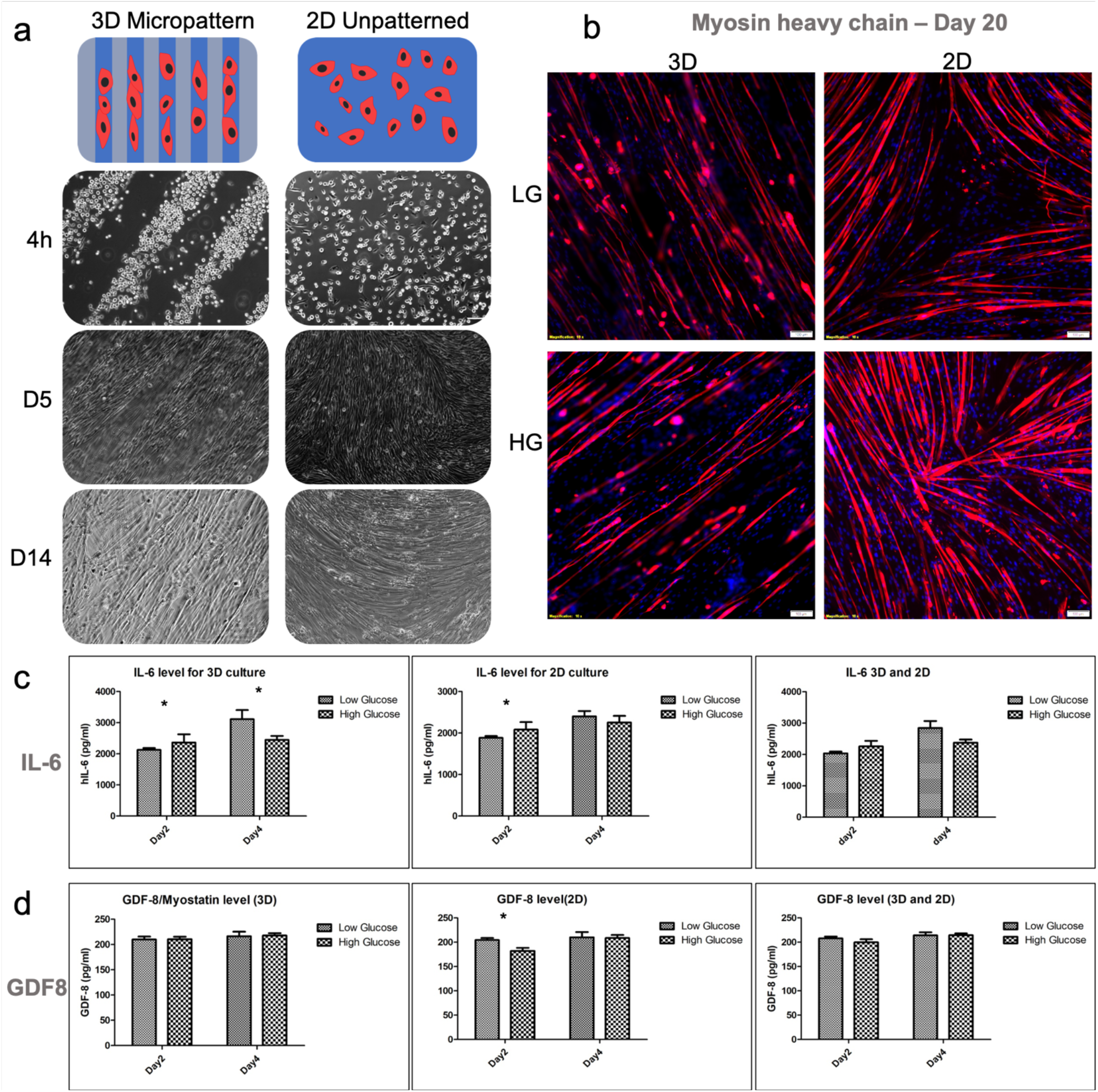
Evaluation of topographical patterning of HA bioinks to induce alignment and fusion of human skeletal muscle progenitor cells. a) 3D micropatterned and 2D unpatterned bioink substrates with human skeletal progenitor cells cultured for 14 days in skeletal muscle differentiation media. Light microscopy images were taken 4 hours, 5 days, and 14 days after cell incorporation. b) Immunofluorescent staining of myosin heavy chain in differentiated cells on 3D micropatterned bioinks versus unpatterned bioinks in low glucose (LG) or high glucose (HG) media. Red – myosin heavy chain; Blue – DAPI. Scale bars – 100 μm. c-d) ELISA-based quantification of c) IL-6 and GDF-8 myostatin in 3D micropatterned bioinks or unpatterned bioinks in low glucose (LG) versus high glucose (HG) media. Statistical significance: * p < 0.05.

### Bioink support of pancreatic islets

Over the past 5 years, we have optimized the environmental conditions that cells and cellular spheroids or organoids experience during the phases of the bioprinting process to the point that viability of printed tissue (and tumor) constructs has largely been a non-issue, demonstrating that our bioinks are sufficiently supporting the cells from a biological, biochemical, and mechanical perspective (*11, 18, 22, 27, 44, 45*). However, pancreatic islets, which are crucial for any diabetes-related studies, have been notably difficult to handle *ex vivo/in vitro* for biofabrication applications due to their inherent fragility during processing (*46*). To address this hurdle, we specifically assessed the viability of human pancreatic islets after having been bioprinted in 4 different bioinks (***Fig. 6a***). These included our HA-Coll bioink, a commercially available bioink (Cellink), and two experimental nanocellulose bioink formulations provided by the Gatenholm laboratory (Chalmers University of Technology, Gothenburg, Sweden). All 4 bioink formulations performed well in terms of printability during the printing process. As evidenced by LIVE/DEAD staining and macro-confocal microscopy images, all bioinks yielded hydrogel constructs with more viable cells than dead cells at day 1. While more dead cells were observed in our HA-Collagen bioink on day 1 compared to the other bioinks, it appeared that long-term viability in the HA-collagen bioink was comparable or better than the other bioinks tested. We noted robust cell viability and maintenance of islet structure in HA-Coll bioinks, which was confirmed by hematoxylin and eosin staining (***Fig. S8a***). In comparison, the 2^nd^ nanocellulose bioink did not support islet physical integrity, and the other bioinks generally resulted in islets of increasingly smaller size over time. Furthermore, on days 1 and 2 following bioprinting, we were able to detect expression of both glucagon and insulin in our HA-Coll-printed islets (***Fig. S8b***), suggesting relevant functionality of the engineered constructs.

**Figure 6.**
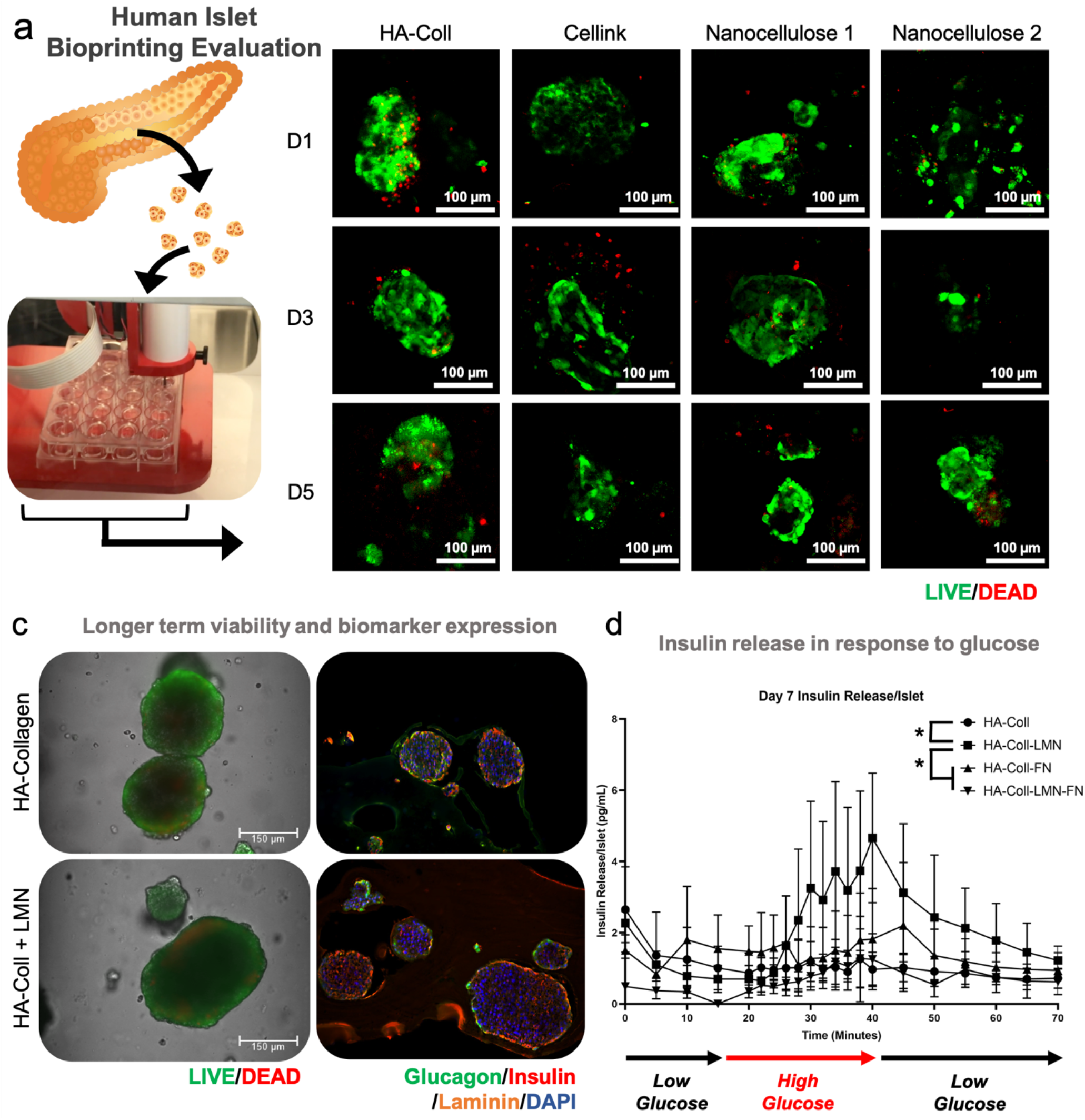
Assessment of human pancreatic islets in bioinks and functional islet characterization. a) Human pancreatic islets were bioprinted in the HA-Coll bioink, the commercially available CELLINK bioink, and two nanocellulose formulations. Relative viability is visualized by LIVE/DEAD staining and macro-confocal imaging. Green – Calcein AM-stained viable cells; Red – Ethidium homodimer-1-stained dead cell nuclei. c) (left column) LIVE/DEAD staining of islets in HA-Coll versus HA-Coll + laminin. (right column) Immunohistochemical analysis of islets in HA-Coll versus HA-Coll + laminin. Green – glucagon; Red – insulin; Orange – laminin; Blue – DAPI. d) Quantification of insulin secretion in response to high glucose conditions by human pancreatic islets in HA-Coll, HA-Coll-LMN, HA-Coll-FN, and HA-Coll-LMN-FN bioinks. Statistical significance: * p < 0.05 between HA-Coll and other experimental groups when considering each data set as a whole Gaussian curve.

These initial studies supported continued experimentation with our HA-Coll bioink in the context of pancreatic islet cultures. Based on established islet biology and histology, it is known that LMN plays an important role as a basement membrane surrounding islets in the pancreas, potentially serving as a stabilizing agent. Based on this understanding, and the modular features of our bioink system, we evaluated the influence of the addition of LMN within the bioink in terms of islet viability and function. LIVE/DEAD assays showed islets that did not vary significantly in terms of overall viability between HA-Coll and HA-Coll+LMN bioinks (***Figure 6c, left side***). However, when assessing presence of glucagon, insulin, and LMN through fluorescent immunohistochemistry, we observed a qualitative difference. Glucagon was expressed in islets in both conditions. LMN presence was increased in the +LMN conditions, as would be expected based on its inclusion in that bioink formulation. However, insulin expression appeared more widespread throughout islets in the +LMN bioinks (***Figure 6c, right side***). While qualitative, this highlighted the need for additional assessment of islets versus bioink ECM composition in a functional assay.

One of the crucial functions of islets is the production of insulin in response to glucose. We prepared islet constructs as described above in our HA-Collagen only bioinks, or with LMN and FN added separately or together. After 7 days of culture in baseline low glucose conditions, an insulin response study was initiated. Following initiation, after approximately 15 minutes, LG media was replaced with HG media for 25 minutes, after which HG media was replaced with LG media. During this 70-minute protocol, media aliquots were removed, frozen, and later assessed for insulin concentration. ***Fig. 6d*** shows insulin release per islet on average. The variability in the experimental groups was quite large and as such direct comparisons between groups at individual timepoints were not statistically significant. However, we noted an overall pro-insulin secretory response only by LMN-containing bioinks. All other bioink formulations showed minor responses to the glucose-stimulated insulin secretion challenge. Instead of considering statistical comparisons at singular time points, we treated each group as a Gaussian curve determined by a nonlinear regression analysis and performed one-way ANOVA analysis of the entire curves against one another. In this statistical analysis, the LMN-containing bioinks did yield a statistically significant increase in insulin production in response to the HG condition.

### Bioink induction of adipose tissue differentiation and construct response to insulin

Adipose tissue plays an outsized role in terms of being involved in more disease states than many other tissues. It is obviously involved in diabetes, but adipose tissue is more dynamic than generally appreciated. Specifically, adipose tissue responds to glucose levels by secreting cytokines that have downstream impacts on other tissues. Adipose tissue biology has implications not only in diabetes, but also cancer and forms of dementias. From a clinical and commercial point of view, adipose tissue is a useful resource that can be harnessed in reconstructive surgeries. Thus, generation of adipose tissue constructs could be valuable both in the realm of microphysiological systems and *in vitro/ex vivo* disease modeling as well as clinical translation for use in human patients.

Initial studies focused on the culture of human preadipocytes in the same 4 bioink formulations described above (HA+Coll (control), +FN, +LMN, and +FN+LMN). Differentiated adipose constructs were imaged under light microscopy to evaluate general cell morphology (***Fig. S9***). HA+Coll, +FN, and +LMN conditions all resulted in compact constructs with preadipocytes that stayed either in a small volume or formed cell aggregates (***Fig. S9a-c***). The HA-Coll+FN+LMN group resulted in a 3D construct that increased in size, and in which cells spread out with a qualitatively healthier morphology (***Fig. S9d***).

Next, the preadipocyte constructs were evaluated for readiness for differentiation into adipose tissue constructs. After 4 days in culture, constructs using the 4 bioink formulations were cultured in preadipocyte media or adipocyte differentiation media for 3 days. The organoids were then maintained in adipocyte nutrition media for 2 weeks for the differentiation process (i.e., mature adipocytes) to be completed. Constructs were then assessed for intracellular lipids by Nile Red fluorescent staining. We specifically decided not to use Oil Red staining based on testing between 2D cultures and 3D spheroids. While 2D cultures provided good visualization of Oil Red staining (***Fig. S9e-f***), 3D spheroids were far more difficult. The stain was only visible in cells that had broken away from the spheroid (***Fig. S9g-h)***. As such, we employed Nile Red fluorescent staining with macro-confocal imaging to better assess differentiation of preadipocytes to adipocytes in the bioink-supported 3D tissue construct format (***Fig. 7a***). In all conditions in which adipose differentiation media was used, constructs exhibited increased Nile Red staining, indicating successful adipogenic differentiation. In the base Coll+HA and +FN conditions without differentiation media, very little Nile Red staining was observed. Remarkably, even without differentiation media, +LMN and +LMN+FN bioinks resulted in similar lipid production in comparison to all cultures using adipose differentiation media, suggesting that the use of these adhesion proteins could drive adipogenic lineage-like commitment on their own. Importantly, when we evaluated expression of a panel of biomarkers via qRT-PCR, we noted that our bioengineered adipose constructs exhibited increased expression of GLUT4, LPL, LEP, LGALS1 compared to PPARG, confirming that the differentiated phenotype observed was in fact accurate (***Fig. 7b***).

**Figure 7.**
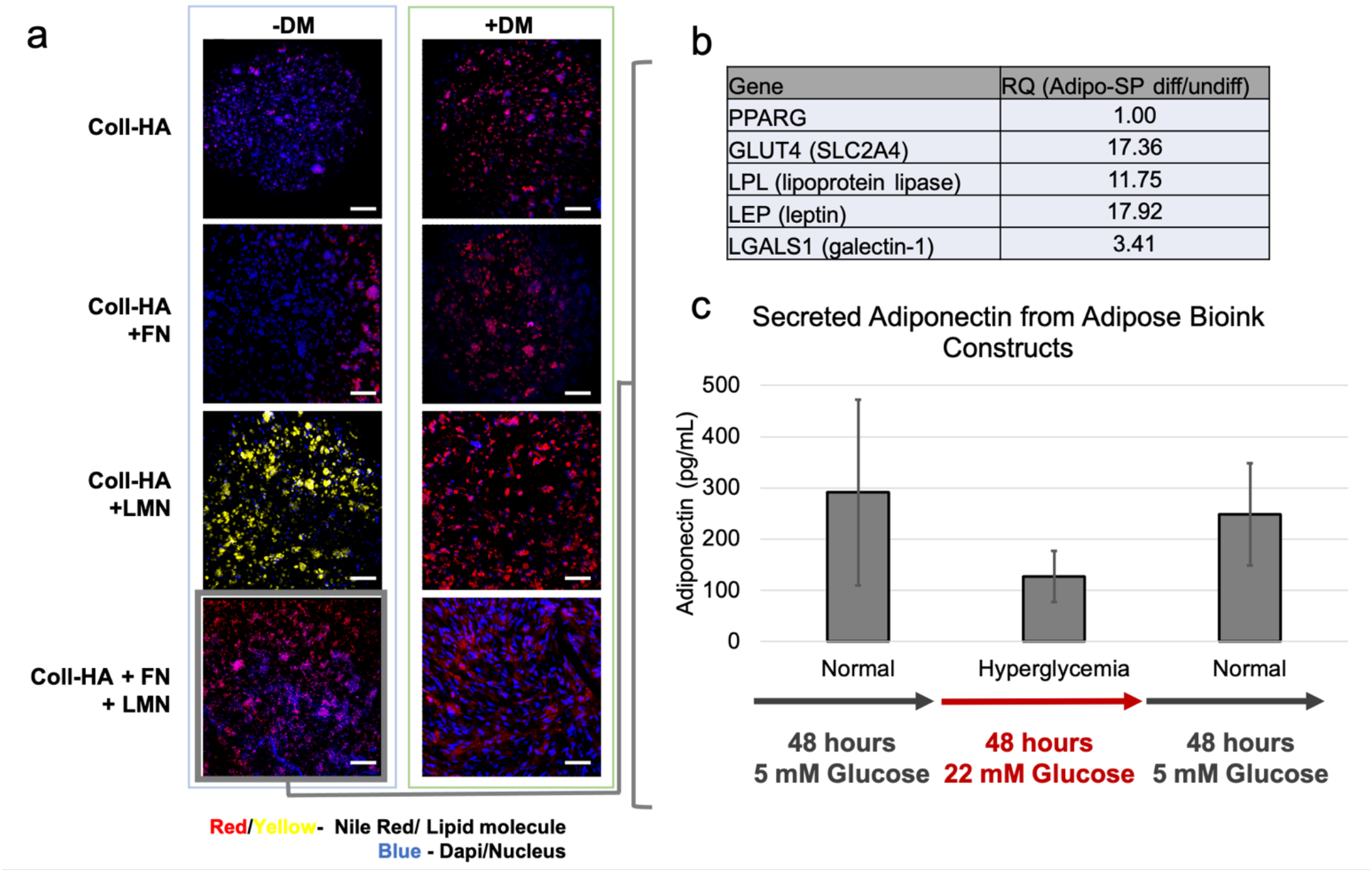
Bioink formulations drive adipose tissue differentiation. a) Preadipocytes encapsulated in HA-Coll, HA-Coll-LMN, HA-Coll-FN, and HA-Coll-LMN-FN bioinks display distinct readiness to differentiate towards adipocytes. With differentiation media (+DM), all constructs stain positively for Nile Red. Without differentiation media (-DM), Coll-HA+LMN and Coll-HA+FN+LMN bioinks drive adipose differentiation. b) qPCR expression in bioink-supported adipose constructs. c) Functional testing of adipose constructs using low glucose to high glucose shift. Adiponectin output is measured by ELISA under normal, high, and normal glucose levels.

Lastly, as a test of functional output, we employed a simplified HG insult assay similar to that used to assess our pancreas islet bioink formulation. As with islets, adipose tissue normally responds to high glucose levels in the blood by decreasing secretion of adiponectin, which is a protein hormone with roles in regulating both glucose levels and fatty acid breakdown (*47*). To assess whether our bioink supported adipose constructs, samples were maintained in 5 mM glucose media (LG) for 48 hours, after which media was replaced with 22 mM glucose media (HG) for 48 hours, and then replaced with LG media once more for 48 hours. At the end of each 48-hour block, media aliquots were obtained and analyzed for adiponectin concentration by ELISA. Interestingly, we observed a reduction in adiponectin from an average of approximately 300 pg/mL under LG conditions to approximately 125 pg/mL under HG conditions (***Fig. 7c***). After returning to LG conditions in the final stage, adiponectin returned to an increased level of approximately 250 pg/mL. While these values were not statistically significant from one another, the trend in this dynamic response to glucose shifts is promising and may be an important function for future studies centered around building an *in vitro* multi-tissue platform to study diabetes.

## Discussion

Bioprinting, and more generally, 3D biofabrication strategies have been long touted as game changing technologies in the regenerative medicine and tissue engineering space. We were promised bioprinted organs as early as 2005, at least by speculation (*48–55*). At this point, “bioink” meant cellular mass (spheroids) (*51, 56*) while “biopaper” referred to the biomaterial in which cells or cell spheroids would be printed into. The discrepancy between bioink and biopaper in the context of bioprinting is no longer, as the term biopaper has largely disappeared, while bioink has come to describe any material that can be deposited from a bioprinter, including both biomaterials and cells, but today leaning more towards the biomaterial component. In these earlier days of bioprinting, we demonstrated that naturally derived ECM-based biomaterials could be harnessed and bioprinted for the first time, driving the concept of a biomaterial-based bioink (*15, 16, 28*). Approximately a decade has passed since these studies, and today bioprinting is much more widely used. However, while the hardware has advanced, development of bioink technologies have lagged. Most laboratories utilize overly simplistic materials to adhere either to native biochemical signals (*e.g.*, Matrigel, which natively is not print-friendly) or highly printable materials that largely ignore crucial ECM signaling (e.g., alginate-gelatin mixtures). In the work we describe here, we sought to approximate and combine the biochemical properties of human tissues with mechanical properties that support printability, while sacrificing neither.

Interesting efforts have been made by many in the bioprinting field to address the so called biofabrication or printability window (i.e., the timeframe to crosslink the bioink between extrusion and deposition) (*57*). Perhaps the most common approach has been to use blends containing gelatin, which serves as a superbly extruding bioink at a semi-cooled state, and is then washed out leaving behind the other post-printing crosslinked component of the bioink (*58*). These efforts employ a secondary material such as alginate, gelatin-methacrylate, HA-methacrylate, and PEGDA – all of which 1) do not contain native cell adherent peptide motifs, with the exception to gelatin, 2) fail to provide a crucial collagen component that cells can reorganize into relevant micro-architectures and aligned fibers, and 3) do not recapitulate the complexity of the native tissue ECM (*21, 59*).

Our approach was built upon a straightforward, yet effective HA and collagen bioink that was printable, supported cell-mediated ECM remodeling (*22*), and could be further modified in terms of printability through the generation of a thixotropic system (*44*). Just as important was devising a strategy to imbue biochemical profiles of tissues, but in a defined manner. In other words, rather than using Matrigel or tissue-derived decellularized ECM materials, we sought to build from the ground up using a discrete set of components. We believe this approach to be more feasible in terms of regulatory hurdles for future regenerative medicine biomanufacturing goals and for development of tissue engineered tools for clinical diagnostics. To address these goals, we first focused on printability.

Naturally derived ECM biomaterials have been notoriously incompatible with most printing platforms with the exception of gelatin, due to its thermoreversible properties. In past studies, we have used chemistry to modify materials such as HA, gelatin, and collagen to generate defined ECM bioinks (*15, 16, 18, 28, 44, 45, 60*). However, these bioinks largely depend on timing of crosslinking or multiple stepwise crosslinking reactions (*18, 19, 28*). In a similar vein to a recent publication that utilized nanoparticles to induce inter-hydrogel forces that resulted in thixotropic mechanical properties (*44*) (i.e., the biomaterial responds to external forces by failure of internal bonds, resulting in dynamic reduction, followed by recovery of elastic modulus) we used catecholamine compounds that linked into our bioinks to further modulate thixotropy and generate bioprintable ECM-based materials. Introduction of catecholamines was originally meant to imbue hydrogen bonding capabilities to generate sticky ECM hydrogels that could potentially serve as surgical adhesives with regenerative properties. However, we realized that by modulating the ratio of covalent to hydrogen bonds within the hydrogels, the extent of thixotropy could be tuned (***Fig. 1h***). As with HA and collagen (or gelatin) bioinks, this strategy could be deployed in most other hydrogels provided they have compatible functional groups for covalent tethering of the catecholamines. Notably, while the studies described herein primarily utilized the CELLINK BIO X and Allevi Allevi2 bioprinters, we also tested these bioinks in additional bioprinting hardware platforms, including the CELLINK Inkredible, the Advanced Solutions Biobot, and the ITOP (Integrated Tissue-Organ Printer) custom platform developed at Wake Forest (***Fig. S10***) (*12*).

In general, we kept the concentrations of thiolated FN and LM in bioink formulations consistent across the board to demonstrate the potency of an approach that focused on simple ECM component changes and subsequent tissue-specific output metrics. Our results logically present several obvious offshoot future studies, in which modulation of concentration of FN or LMN as the primary independent variables in a more nuanced fashion could be investigated. Indeed, our team is currently assessing the influences of subtle shifts in these ECM components within our ECM bioinks on tumor and immune cell 3D migration in the context of bioengineered *in vitro* and *ex vivo* cancer models (*61, 62*). One important consideration we assessed was whether or not FN and LMN need to be immobilized to the bioink components, or whether soluble FN and LMN, encapsulated within the bioink, but not immobilized, was important. As we demonstrate in two examples (***Fig. S2***), covalent bonding of the adhesion proteins within the bioink was crucial to facilitating cell-ECM interactions and attachment. Without covalently bound proteins, cells adhered to one another rather than the ECM, forming aggregates which would be washed away in routine staining protocols. Based on these data, our thiolated versions of these proteins were employed in all subsequent tissue-specific studies.

In most of our tissue-specific studies, we observed a clear benefit from bioink customization via addition of thiolated FN, LMN, or both. This included liver constructs, both in terms of viability and function, neural cultures, including both neuron-like cells and the BBB construct, skeletal muscle, and adipose constructs, in which the bioink formulation drove differentiation towards adipose tissue, without the need for differentiation media. Pancreatic islet cultures were promising. The HA-Coll bioink appeared to maintain islet integrity through the bioprinting process more readily than other bioinks tested. Moreover, the addition of thiolated LMN appeared to increase insulin expression, as noted by immunofluorescent staining and insulin release studies. However, admittedly, the insulin release studies were not statistically significant, but showed a clear trend, both on day 1 and day 7 in culture, setting the stage for a future study focused specifically on islets and further bioink formulation modulation. One notable observation was that only the LMN-containing bioink showed measurable increases in insulin secretion in response to high glucose. FN-containing formulations, and even FN+LMN-containing formulations did not. This is in line with understood physiology of the pancreas, where LMN is present around the circumference of healthy islets, which supports glucose response and insulin release (*63, 64*), while increased FN concentrations correlates with several disease states such as cancer (*65*). Interestingly, skeletal muscle constructs were not particularly dependent on ECM composition, but rather were highly dependent on architecture that facilitated cellular alignment. This also warrants future study, as generating a fully 3D hydrogel-based construct in which “topography” is maintained through the 3D space is a challenge. Internal ECM fiber alignment (e.g., collagen) may be a solution for these models.

Our goal is to apply this bioink platform in two distinct areas of tissue engineering. First, and most immediate, is within the space of microphysiological systems, or organoids and organ-on-a-chip platforms. Our team has previously demonstrated the capability to generate multi-tissue systems containing organoids and micro-tissue constructs in which we show inter-tissue interactions that cannot be observed in single-tissue models (*11, 33, 34, 37, 66*). With the system described herein, we can use a set of 4 ECM base components to support a variety of tissue types with high *in vitro* and/or *ex vivo* functionality. Therefore, an immediate interest is in developing a more functional diabetic “body-on-a-chip” platform. Efforts have been made in this area, but have largely been limited to liver and islet systems using spheroids, 2D cultures, a mix of these formats, and in one example a muscle component (*67–69*). While these are important advances, they generally fail to account for the role of ECM, and as such there is significant room to further develop this technology area. Additionally, we are actively deploying these bioink formulations in the realms of other disease models, with a focus in modeling cancer, cancer metastasis, and the progression of the pre-metastatic niche. However, the versatility of these materials is also being used for other disease models, including glaucoma for example (*70*). Furthermore, we have used the base materials of this platform, and earlier iterations of our bioink systems, for creating a patient-derived tumor organoid platform with which we have performed chemotherapy and immunotherapy screening studies across a variety of cancer types (*45, 61, 62, 71–74*). There is an incredible opportunity to deploy such systems that are constructed using defined, FDA-approved materials in clinical settings such as diagnostic drug screening assays to aid in optimizing treatments for human patients (*61, 75*).

Second, the well-defined nature of our bioink technology can be leveraged towards future regenerative medicine and tissue engineering applications in the clinical and commercial spaces. Regenerative medicine-based biomanufacturing is a realm that is growing, albeit with many regulatory hurdles (*6, 7*). By using a bioprinting-friendly biomaterial platform of defined, xeno-free composition, we provide a potential toolkit to translate tissue engineering applications based on ECM formulations with far more nuance than simply gelatin-methacrylate, alginate, methacrylated hyaluronan, and PEG-based hydrogels. These oft used bioink materials can be harnessed effectively for printability but fail to encompass even the bare range of ECM signals that are required for advanced tissue biofabrication, support, and maturation (*14*). Our platform is not without limitations. It is based on only 4 primary ECM components; yet, this is still more encompassing of ECM biology than other defined bioinks. Our bottom-up approach based on corresponding chemistries of each component can be further expanded. For example, we are exploring substitution of whole adhesion protein modifications to minimization to the adhesion peptide sequences of FN and LMN with cysteines providing thiols for crosslinking. We are also evaluating modification of additional collagens (III and IV are of particular interest) for direct crosslinking into our platform.

In summary, our approach offers a defined, regulatory-ready platform that provides more nuanced control over tissue-specific support in tissue engineering applications. It is xeno-free, and more importantly not derived from tumorigenic animal tissue, thus setting it up for future translational applications requiring regulatory approval. While being well-defined, this system allows for user customization and support of the multiple tissue types described herein, as well as ongoing studies in our laboratory in biofabricating advanced patient-derived tumor models and blood malignancies such as myeloma. With our continued work with this ECM hydrogel platform we anticipate deployment in both diagnostic and drug screening applications as well as more complex clinically oriented tissue engineering of tissues and one day organs for transplantation in human patients.

## Materials and Methods

### HA and Collagen base bioink preparation

Bioink base formulations were prepared by combining methacrylated collagen (Coll, Advanced Biomatrix, San Diego, CA) and thiolated HA (HA, formerly ESI BIO, Alameda, CA; now Advanced Biomatrix) as we have described previously (*22*). Coll was prepared at 6 mg/mL per manufacturer’s instructions excluding the provided photoinitiator. Prior to use with HA, Coll was neutralized using manufacturer provided neutralization solution at 85 μL of solution per milliliter of collagen. HA was prepared at 2 mg/mL by re-suspending Heprasil (heparinized and thiolated HA) in 1 mL deionized water with 0.1% w/v photoinitiator (2-Hydroxy-4′-(2-hydroxyethoxy)-2-methylpropiophenone, Irgacure D-2959; MilliporeSigma, St. Louis, MO). In general, the ratio of 3 parts Coll to 1 part HA by volume was used. Following mixing, hydrogels – with or without cells – were then deposited as needed by the bioprinter, as described below, or in some cases manually for initial tests, or in subsequent stages of the study with additional component inclusion based on the particular tissue type. Deposited bioink volumes were crosslinked with UV light (365 nm, 4.66 W/cm^2^/second Dymax BlueWave 75, Torrington, CT) for about 2 seconds each to crosslink the polymer networks in place.

### Catecholamine synthesis for thixotropic bioinks

For catecholaminolkyne (2-(3,4-dihydroxyphenyl)-N-(2-proynl)acetyl [catecholaminoalkyne]) synthesis, first 3,4-dihydroxybenzaldehyde (500.0 mg, 3.62 mmol, 1.0 eq, MilliporeSigma) was suspended in 15 mL of dry dichloroethane (DCE, MilliporeSigma). Subsequently, propargylamine (347 μL, 5.43 mmol, 1.5 eq) was added dropwise and stirred for 30 minutes at room temperature under inert conditions. Sodium triacetoxyborohydride (2.148 g, 10 mmol, 2.8 eq, MilliporeSigma) was added and allowed to react for 24 hours. No purification methods followed and the synthesis resulted in an isolated yield of 98%. The synthesis of the compound was confirmed with ^1^H-NMR and ^13^C-NMR in a recent publication (*26*). See **Figure 1d** for chemical structure.

For catecholamidoalkyne (2-(3,4-dihydroxyphenyl-N-(2-propynyl)acetamide) [catecholamidoalkyne]) synthesis, 3,4-dihydroxybenzoic acid (1.00 g, 6.49 mmol, 1.0 eq, MilliporeSigma) was first dissolved in 30 mL of acetonitrile and was followed by the dropwise addition of propargylamine (831.3 μL, 12.9 mmol, 2 eq, MilliporeSigma). 1-Ethyl-3-(3-dimethylaminopropyl)carbodiimide (EDCI, 1.21 g, 7.79 mmol, 1.2 eq, MilliporeSigma) was slowly added and the resulting reaction was refluxed at 70°C for 3 hours. Once completed, the compound was purified by column chromatography (silica gel, solvent gradient of 50%, 80% and 100% ethyl acetate [EtOAc] in pentane), with an isolated yield of 75%. The synthesis of the compound was confirmed with ^1^H-NMR and ^13^C-NMR in a recent publication (*26*). See **Figure 1e** for chemical structure. More detailed description and analyses of these compounds have been recently published (*26*).

### Rheological mechanical testing of thixotropic properties

For initial testing, a gelatin variety of our bioink, based on the HyStem hydrogel kit, was employed to more directly control the ratios of covalent versus hydrogen bonding. To a 2 mL centrifuge tube, 200 μL of 1% w/v HA in water, 200 μL of 1% thiolated gelatin in water, 50 μL of 2% w/v polyethylene glycol diacrylate in water, and 50 μL of a modular catecholamine or amide solutions (0.25%-1.5% w/v) were added and mixed by gently pipetting the solution up and down to avoid creating bubbles. For control experiments the modular catecholamide or -amine solutions were replaced with an equivalent volume of sterile deionized water. For rheology or cell culture studies, the reaction mixture was placed into wells of an appropiate size and exposed to UV radiation (365 nm, 4.66 W/cm^2^/second Dymax BlueWave) for about 2 seconds each. See **Figure 1f-g** for crosslinking schematic and chemical diagram.

To assess the thixotropic behavior of the catecholamine-modified hydrogel, a multistep hysteresis loop test was performed. A Discovery HR-2 Rheometer (TA Instruments, New Castle, Delaware) with an 8 mm geometry was used to collect the rheological data. Volumes of 200 μL of the reaction mixtures were transferred into 12 mm diameter × 5 mm depth custom fabricated PDMS wells. The PDMS well containing the reaction mixture was then exposed to UV light (365 nm, 4.66 W/cm^2^/second for about 2 seconds from 1 cm). To ensure standard conditions across all experiments the geometry was lowered into the gels until a calibration normal force of 0.4 N was achieved. The hysteresis loop test was performed after an initial sample conditioning (calibration) step, after which an increasing logarithmic strain sweep (0.2 to 1.0%) was performed. This was followed immediately with a regeneration pause at a constant oscillation amplitude, and lastly a decreasing logarithmic strain sweep (1.0% – 0.2%). To present the hysteresis loop data, the storage (elastic) modulus, ***G’*** was plotted against the oscillation strain percentage.

### LIVE/DEAD viability/cytotoxicity staining and ATP activity quantification

Throughout the study, biocompatibility of the different gel formulations or tissue construct types was assessed using a standard LIVE/DEAD cell viability/cytotoxicity kit (Thermo Fisher, Waltham, MA) according to the manufacturer’s recommendation. Briefly, the cell membranes of viable cells are stained with calcein AM and have a green fluorescence emission. Dead cells are identified by ethidium homodimer-stained cell nuclei with a red fluorescence emission. Stained organoids were imaged using macro-confocal microscopy (Leica TCS LSI, Leica Microsystems, Buffalo Grove, IL).

Proliferation of cells within the different constructs was generally evaluated on days 4, 7, and 10 (n = 3) by quantification of mitochondrial metabolism with a CellTiter 96® Aqueous One Solution Cell Proliferation Assay kit, also known as the MTS assay, (Promega, Madison, WI) according to the manufacturer’s protocol. Absorbance was quantified on a Spectramax M5 plate reader (Molecular Devices, Sunnyvale, CA) at 490 nm.

### Protein thiolation methods, cell adherence validation of protein integration, cell line viability assessment, and cell phenotype evaluation

Lyophilized human FN and LMN protein (purified from plasma, MilliporeSigma) was dissolved in 1X Dulbecco’s Phosphate-Buffered Saline (DPBS) at pH 7.4 containing 10 mM EDTA for a final concentration of 1 mg/mL. N-Succinimidyl-S-acetylthioacetate (SATA; MilliporeSigma) was dissolved in dimethyl sulfoxide (DMSO; MilliporeSigma) for a concentration of 65 mM. Fifty mL were added immediately to the 1 mg/mL FN or LMN solutions and allowed to react for 30 minutes at room temperature. The reaction mixture was dialyzed against 1X DPBS pH 7.4 containing 1 mM EDTA for 4-6 hours to remove unreacted SATA. After dialysis, 500 mL of a 0.5 M hydroxylamine in 1X DPBS at pH 7.4 containing 25 mM ethylenediaminetetraacetic acid (EDTA) was added to the reaction mixture and left undisturbed for 2 hours at room temperature. Then, the reaction mixture was dialyzed against 1X DPBS at pH 7.4 containing 1 mM EDTA overnight and the dialysis buffer was changed every 6-8 hours. Lastly, a buffer exchange with deionized water was performed to remove salts prior to lyophilizing. After lyophilizing, the thiolated FN was stored at −20°C and reconstituted in 1X DPBS pH 7.4 immediately prior to use.

As functional validation of thiolated adhesion protein integration, thiolated FN was employed as a test case. First, an adherent colorectal cancer tumor cell line, HCT-116 (ATCC.org, Manassas, VA), was plated on thiolated HA gels crosslinked with PEGDA. Second, thiolated HA, thiolated FN, and PEGDA were crosslinked together to form FN-containing hydrogels. Third, thiolated HA, thiolated gelatin, and PEGDA was prepared and crosslinked. HCT-116 cells were plated on these substrates at a seeding density of 20,000 cells per well in a 24-well plate. Following 24 hours, each well was assessed for cell attachment by brightfield imaging. Subsequently, wells were washed with PBS, and stained with phalloidin 594 (Abcam) and DAPI, and imaged by fluorescent microscopy to determine whether cells adhered and spread out on the hydrogel substrates. This was then repeated using human umbilical cord endothelial cells (HUVEC, Lonza, Walkersville, MD) on HA+gelatin+PEGDA, HA+PEGDA with FN added to the media supernatant, HA+unthiolated-FN+PEGDA, and HA+FN+PEGDA hydrogels. Brightfield microscopy was used to assess cell-cell aggregation versus cell adherence onto the hydrogel surface.

Initial effects of including thiolated proteins on cell viability were assessed using a panel of commonly used cell lines obtained from ATCC.org. Following expansion on tissue-treated dishes with Dulbecco’s Minimum Essential Medium (DMEM), the cells were trypsinized with 0.05% of trypsin (Hyclone, Logan, UT) and counted. Tissue constructs of each model cell line (HepG2 [human hepatoma], U-87 MG [human glioblastoma], and HCT-116 [human colorectal carcinoma]) were created using 50,000 cells of the respective cell type and suspending them in either 10 μl of control HA-Gel hydrogel formulation (HA, gelatin, and PEGDA) or in 10 μl of formulation of HA-Coll gel (collagen, HA, and FN) to verify that the collagen and FN containing hydrogel showed equally sufficient viability as the commercially available formulation. We chose this cell density for organoid/cell construct formation based on the results obtained from 3D organoid research performed in our lab (*33, 45, 66, 76, 77*). The hydrogel precursor-cell mixtures were pipetted into wells of a 48-well plate (Corning, NY) that was pre-coated with 200 μl of PDMS (Sylgard 184 elastomer kit, Dow Corning, Midland, MI) prepared according to manufacturer’s protocol to create a hydrophobic surface. The cell constructs were then crosslinked with UV light (365 nm, 4.66 W/cm^2^/second for about 2 seconds from 1 cm, Dymax BlueWave 75). The wells were filled with 500 μL of media, which was replenished on day 4 and day 7 of the experiment. The cell constructs were maintained in culture for 10 days. LIVE/DEAD staining and imaging and ATP activity quantification was performed, as described above, on days 4, 7, and 10.

Addition of thiolated LMN and FN were then assessed for their ability to directly drive cell phenotype in terms of epithelial versus mesenchymal morphology. HA-Coll hydrogels were formulated as described above, supplemented with either LMN only, FN only, or both LMN and FN. These hydrogels were used to encapsulate LX2 stellate cells (previously expanded on tissue culture plastic using DMEM 10% FBS, 1% pen/strep, and 1% L-glutamate) at a cell density of 10 million cells/mL. Tissue constructs were maintained for 7 days, after which they were stained with phalloidin-488 (Abcam) and DAPI. Stained constructs were imaged using macro-confocal microscopy (Leica TCS LSI, Leica Microsystems).

### Hydrogel characterization: Rheology, porosity, and swelling

Hydrogel mechanical properties were assessed by rheology, comparing the three hydrogel formulations: HA-Coll, HA-Coll-FN, and control HA-Gel, thereby assessing whether or not the addition of a thiolated adhesion protein significantly influenced material properties. Once all components of each hydrogel were mixed and crosslinked, the hydrogels were analyzed using a TA instruments DHR-2 rheometer (TA Instruments) with a 25 mm plate and 25 mm 2 ° cone system. The rheology test was comprised of a strain sweep from 1% to 1000% shear strain (γ), 10 points per decade, in a logarithmic sweep, performed at a frequency of 1 Hz. Reported were shear elastic moduli ***G’*** and shear loss moduli ***G”***.

To evaluate porosity, HA-Coll, HA-Coll-FN, and HA-Gel hydrogels were frozen and lyophilized. Resulting freeze-dried hydrogels were cut to expose the inner meshwork surface and gold sputtered, after which microstructures in the cross-sections were imaged by scanning electron microscopy using a FlexSEM 1000 (Hitachi, Ltd., Tokyo, Japan). Pore size was assessed using ImageJ software (National Institutes of Health) after image calibration. Average pore sizes for each experimental group were determined based on 15 measured pores.

Swelling behavior was determined by standard swelling assays as previously described (*29*). First, a 0.1 mL aliquot of each hydrogel type was formed in the bottom of replicate glass vials and the vials weighed. Then, 1 mL of DI water was placed over of each gel, and the vials were placed into an incubator at 37°C. In this case, swelling of the hydrogels occurred uniaxially toward the upper direction of the vials. After 24 hours, the vials were weighed after carefully removing the DI water. The mass of the both the initially formed, and swollen hydrogels was calculated by subtracting the mass of the vial from the total mass. The swelling ratio was defined as a ratio of the mass of the swollen hydrogel divided by the mass of the initial hydrogel.

### Bioprinting methodology

The respective bioink formulations were prepared using methacrylated Coll I 6 mg/mL and thiolated HA 2 mg/mL in a 3:1 ratio, modulated as described above for print testing and thixotropy assessment. This base bioink was then mixed with thiolated LMN and thiolated FN depending on tissue type as described below in each respective section, after which cell components were resuspended in the bioink precursors. The hydrogel–cell mixtures were transferred into 10 mL syringe print cartridges with 23-gauge needles and loaded into the extrusion printhead and kept at 4C° through temperature control, until ready to print. Following printer calibration (Allevi Allevi2, Allevi, Philadelphia, PA; or Cellink BIO X, Cellink, Boston, MA), custom written 24- or 48-well plate G-code files were used to direct the printer to extrude 10 μL of hydrogel bioink–cell mixtures into the middle of the well at 3 PSI pressure and a speed of 20 mm/s. Following the printing process, all printed organoids/tissue constructs were individually irradiated by UV light (365 nm, 4.66 W/cm^2^/second for about 2 seconds from 1 cm).

### Liver construct biofabrication and basic liver functional characterization

Cryopreserved primary human hepatocytes (Lonza) were thawed according to manufacturer’s protocol and the cells were directly used in experiments. LX2 hepatic stellate cells were trypsinized with 0.05% of trypsin (Hyclone) and counted. Forty thousand human hepatocytes and 10,000 LX2 hepatic stellate cells were suspended in 10 μl of HA-Coll or HA-Coll-FN for the formation of 3D liver constructs. The organoids were engineered in an identical method to cell line organoid formation described above, except deposited in 24-well plates (Corning). The wells were filled with 2 mL of Hepatocyte Culture Media (HCM, Triangle Research Labs), which was replenished after collecting media aliquots for further analysis on day 4, 7 and 10 of the experiment. The organoids were cultured for 14 days.

Cell viability was determined by LIVE/DEAD assay on day 7 and 14 using representative images and by quantification of biomarkers from media collected on days 4, 7, 10, and 14 (n = 3). The secreted levels of albumin and urea of the liver organoids were quantified from media collected on days 4, 7, 10, and 14 (n = 3) using a Human Albumin ELISA Kit (Abcam; Boston, MA. ab108788) and Urea Assay Kit (Abcam, ab83362), respectively. Absorbance was read on a Spectramax M5 plate reader (Molecular Devices) at 450 nm for albumin and at 430 nm for urea.

### Liver construct toxicity screening

The hepatoxic drugs bromfenac sodium, troglitazone and tienilic acid, recalled by the FDA, were chosen for this study.(*31, 32*) All three compounds were purchased from MillporeSigma. Drugs were dissolved in DMSO for stock concentrations of 100 mM. Concentrations of 1 μM, 10 μm, 100 μM, and 1000 μM were used for bromfenac sodium and tienilic acid. Concentrations of 1 μm, 10 μM, and 100 μM were used for troglitazone. The drug toxicity study was conducted for 48 hours in a 24 well plate at 37ºC with 5% CO_2_; each well in the plate containing one liver organoid in 2 mL of media (control) or media with drug dissolved at the concentrations described above (n ≥ 6). The response of the liver organoids was assessed by quantifying relative ATP activity using the Cell-Titer Glo Luminescent Cell Viability assay (Promega) prepared according to manufacturer’s protocol. The results were quantified using the Veritas Microplate Luminometer setup according to manufacturer’s instructions (Turner BioSystems, Promega). Each condition was also subjected to LIVE/DEAD staining (Thermo Fisher). Stained organoids were imaged using macro-confocal microscopy (Leica TCS LSI, Leica Microsystems).

The environmental toxins glyphosate, mercury chloride, and lead chloride were purchased from MilliporeSigma. Drugs were dissolved in diH_2_O for stock concentrations of 50 mM (glyphosate), 10 mM (mercury chloride), and 20 mM (lead chloride). Concentrations of 1 mM, 5 mM, 10 mM, and 20 mM were used for glyphosate. Concentrations of 10 μM, 20 μM, 50 μM, and 100 μM were used for mercury chloride. Concentrations of 1 mM, 2 mM, 5 mM, and 10 mM were used for lead chloride. The environmental toxin toxicity study was conducted for 48 hours (n =7) and liver organoids were assessed for viability with ATP assay and LIVE/DEAD assay as described above.

Acetyl-para-aminophenol (APAP) and N-Acetyl-L-cysteine (NAC), used to treat APAP overdose, were purchased from MilliporeSigma. Drugs were dissolved in HCM media to reach the desired concentrations. On day 7 in culture, liver organoids were treated with no drug, 1 mM APAP, 10 mM APAP, and simultaneously both with 10 mM APAP and 20 mM NAC, respectively (n = 4). Viability of the liver organoids was assessed using LIVE/DEAD assay on day 14; albumin and alpha GST production were quantified from media collected on days 4, 7, 10, and 14. Albumin concentrations were determined as described above. Alpha-Glutathione s-transferase (alpha-GST) is secreted by liver cells upon cell death. The secreted levels of alpha-GST by the organoids were quantified using a GST alpha assay kit (GS41, Oxford Biomedical Research; Rochester Hills, MI). Absorbance was read on a Spectramax M5 plate reader (Molecular Devices) at 450 nm for alpha-GST.

### Neuronal construct biofabrication

Thiolated HA (Heprasil, Advanced Biomatrix), thiolated gelatin (Gelin, Advanced Biomatrix), linear 3400 kDa PEGDA crosslinker (Extralink, Advanced Biomatrix) and 4-arm 10,000 kDa PEGDA crosslinker (Creative PEGWorks) were all reconstituted in 0.1% photoinitiatior (Irgacure D-2959, Sigma Aldrich) at 1% w/v. To create low stiffness hydrogels (200-400 Pa), HA, gelatin, and 2-arm crosslinker were combined 2:2:1 by volume. To create medium stiffness hydrogels (1000 Pa), HA, gelatin, 2-arm crosslinker, and 4-arm crosslinker were combined 2:2:0.5:0.5 by volume. To create high stiffness hydrogels (2000 Pa), HA, gelatin, and 4-arm crosslinker were combined 2:2:1 by volume (*36*). Thiolated FN, LMN, or a combination of both, were added to a final concentration of 0.25 mg/mL. To fabricate neuronal cell constructs, human bone marrow MSCs were cultured in MSC growth medium (PromoCell). The cells were passaged using 0.5% trypsin. MSCs were suspended in 10uL of hydrogel at a concentration of 5×10^6^ cells/mL. These 3D constructs were grown in MSC Neurogenic Differentiation Medium (PromoCell) for 10 days before analysis. Astrocyte-containing hydrogels were fabricated in a similar manner by suspending primary human astrocytes (ScienCell) at a concentration of 5×10^6^ cells/mL in 10uL of the low stiffness hydrogel with the addition of FN and/or LMN.

For the BBB model, brain endothelial cells (BECs) were derived from the IMR(90)-4 (WiCell) human induced pluripotent stem cell line using a previously described protocol.(*78*) Briefly, iPSCs were cultured in mTeSR-1 medium on Matrigel coated plates (Stem Cell Technologies) and passaged into a single cell suspension using Accutase (ThermoFisher). Cells were plated at a concentration of 150,000 cells per 6-well in mTeSR-1 with 10 uM Y27632 (BD Biosciences). The next day, media was changed to Essential 6 media (ThermoFisher) and was replaced daily for 4 days. On day 4, media was changed to human endothelial serum free medium (hESFM, ThermoFisher) supplemented with N2, bFGF, and retinoic acid. On day 6, cells were passaged with Accutase and seeded onto the BBB constructs. The 3D compartment of BBB model underlying the BECs utilized the low stiffness hydrogel with 2-arm PEGDA crosslinker. Primary human astrocytes were grown on the bottom of transwell inserts (24-well, 0.1 um pore size, PET, Fisher Scientific) which were coated with poly-L-lysine (20 ug/mL). Primary human brain microvascular pericytes (ScienCell) were suspended in hydrogels at a concentration of 5×10^6^ cells/mL. The apical surface of the transwell membrane was coated with 50 uL of the crosslinked pericyte hydrogel. Brain endothelial cells were seeded on top of the hydrogel at a concentration of 1×10^6^ cells/cm^2^. BBB constructs were maintained with supplemented hESFM media in the apical chamber and astrocyte medium (ScienCell) and pericyte medium (ScienCell) at a 1:1 ratio in the basolateral chamber. BEC medium was changed to hESFM supplemented with only N2 the day after seeding them in the transwells.

### Neuronal construct functional testing

Cell viability and morphology was qualitatively assessed using a LIVE/DEAD assay as previously described. Trans endothelial electrical resistance (TEER) of BBB constructs was measured using the Epithelial Volt/Ohm Meter (World Precision Instruments, Sarasota, FL). Permeability of the BBB construct was measured by quantifying 5 kDa FITC-dextran diffusion across the barrier. 500 μL of media containing 1 mg/mL FITC-dextran was added to the apical chamber. After 1 h, 200 μL of medium was removed from the basolateral chamber and fluorescence was measured. Values were compared to transwell constructs with hydrogels but no BECs seeded on top. Immunostaining of BBB constructs for tight junction protein ZO-1 was conducted using ZO-1 mouse monoclonal antibody conjugated with Alexa 488 (ThermoFisher). Hydrogels were removed from the transwell insert using a biopsy punch, transferred to a glass slide and imaged with a confocal microscope.

### Skeletal muscle cell culture

C2C12 cells (ATCC, CRL-1772TM) were cultured with high glucose DMEM supplemented with 10% FBS and 1% penicillin-streptomycin (growth media – GM). For spheroid formation, C2C12 cells were seeded into Corning® Costar® Ultra-Low Attachment 96-well plate (MilliporeSigma) at 2000 cells per 100 μl per well in spheroid formation media, which included 20% FBS and 20 μg/ml rat tail collagen I (Fisher Scientific, 354236). Three days after cell seeding, 100 μl per well GM was added. Media was changed every other day by the removal of 100 μl media and the addition of 100 μl per well fresh media. To induce myogenic differentiation, GM was replaced with differentiation media (DM), containing 2% horse serum and 1% penicillin-streptomycin in DMEM. Spheroids were collected and fixed for paraffin embedding and H&E staining. For C2C12 cell topographical patterning experiments, cells were seeded at 5 × 10^4^/well in GM. After 3 days, DM was added to the culture and replaced every other day to induce myogenic differentiation. Myosin heavy chain (DSHB, MF20) immunofluorescence staining was performed to demonstrate cell fusion.

Primary human muscle progenitor cells (hMPCs) were seeded at 1.5 × 10^5^/well and cultured in low glucose (1 g/ml) DMEM supplemented with 5ng/ml bFGF, 20% FBS, 10% horse serum, 1% chicken embryonic extract and 1% penicillin-streptomycin for 6 days. To induce differentiation, media was changed to low glucose DMEM with 10% horse serum for 24 hours and then fed with differentiation media (LG DMEM with 2% horse serum) every other day. hMPCs were differentiated for 5 days prior to the start of the glucose induction experiment. Cells were cultured in either low glucose (1 g/ml) or high glucose (4.5 g/ml) DM. Media was collected at days 0, 2, 4, and 6. At day 6, hMPCs were fixed for MHC immunofluorescent staining. ELISA assays for hIL-6 (R&D Systems, D6050) and GDF-8/myostatin (R&D Systems, DGDF80) were performed using media collected at days 2 and 4.

### Micropatterned bioink skeletal muscle constructs

Micropatterned silicone stamps were fabricated following an in-house soft lithography technique using PDMS (10:1 Sylgard 184 silicone elastomer and curing agent, respectively, Dow Corning, Midland, MI, United States). A 1 cm in length, 200 μm width and 100 μm spacing line-array was design in AutCAD (AutoDesk Mill Valley, CA) in a 2D layout. One layer of double-sided adhesive film (DSF; 140 μm thickness, part number 3M9495MPF, Strouse, MD USA) was pressed on a glass slide to reach the desired pattern heigh of 100 μm. The film was laser cut (Full Spectrum Laser H-series, NV), leaving a negative mold of a 24-line pattern. A permanent resin mold was generated by replication from a PDMS of the DSF device. Epoxy-based negative photoresist SU8 2050 (MicroChem, Westborough, MA) was spin coated onto a glass slide at a height of 100 μm and the positive features of the PDMS replica was pressed onto the SU8. The PDMS/SU-8 arrangement was placed under UV light at 850 mW for 3 min at a 5 mm distance from light source. Positive-PDMS stamps used were replicas derived from the SU8 mold.

Micromolded hydrogels comprised of HA, thiolated gelatin, and PEGDA (Advanced BioMatrix) were made with a PDMS stamp, UV crosslinked on 12 mm diameter round coverslips, and transferred into 24-well plate wells. Generally, 40 μl hydrogel per coverslip was covered with the PDMS stamp and UV crosslinked (365 nm, 4.66 W/cm^2^/second Dymax BlueWave 75) for 2 seconds. After carefully removing the hydrogel from the PDMS stamp, the hydrogel was UV crosslinked for another 2 seconds. Coverslips were transferred into 24 well plate and rinsed with PBS to wash out any uncrosslinked hydrogel.

### Pancreatic islet bioprinting bioink testing and evaluation

The bioprinted pancreatic constructs have an estimated concentration of 70 IEQ (islet equivalency) per 10 μL of hydrogel. Constructs were bioprinted (Allevi2, Allevi) into tissue-treated 24-well plates. Briefly, approximately ~3000 islets/mL were mixed into the bioink. The pancreatic bioink formulation was prepared by combining Coll (6 mg/mL) and HA (2 mg/mL) in a 3:1 ratio. This base bioink was mixed in a 4:1 ratio with a solution of thiolated LMN (5 μg/mL) and thiolated FN (1 mg/mL). The hydrogel–islet mixture was transferred into a 10 mL syringe print cartridge with a 23-gauge needle and loaded into the primary printhead and kept at 4C° through temperature control, until ready to print. Following printer calibration, the proprietary 24-well plate G-code file was employed to extrude 10 μL of hydrogel–islet mixture into the middle of the well at a 3 PSI pressure and a speed of 20 mm/s. Following the printing process, all printed organoids were individually irradiated by UV light (intensity of 1W/cm^2^, dosage of 623 mJ/cm^2^, 10mm distance, for 0.25 seconds). Our HA-Coll bioink was compared to a commercially available bioink (Cellink), and two experimental nanocellulose bioink formulations provided by the Gatenholm laboratory (Chalmers University of Technology, Gothenburg, Sweden). These alternative bioinks were printed under the same conditions. Viability was evaluated by LIVE/DEAD staining and confocal imaging on days 1, 3, and 5 following printing.

### Islet insulin response testing

Pancreatic islet experimentation was carried out to compare hydrogel bioink formulations and their ability to support islets over time, with a focus on the influence of covalently bound FN and LMN. Bioink formulations employed were HA-Coll, HA-Coll+LMN, HA-Coll+FN, and HA-Coll+LMN+FN. In addition, unencapsulated islets were used as a control. The base bioink formulation was used as described above. LMN and FN were added at 5 μg/mL. Human pancreatic islets were purchased from Prodo Laboratories (Aliso Viejo, CA) and allowed to rest in non-adherent tissue culture plastic for two days, then placed into 20 μL hydrogels at 150 islets each. Hydrogels were maintained for 1, 4, or 7 days. At each time point, n=3 hydrogels were fixed and processed for immunohistochemistry and n=3 hydrogels were placed under glucose stress (low glucose (3.3 mM), high glucose (16.7 mM), low glucose, and KCl) and the supernatant was collected after each condition, and insulin was quantified by ELISA (Abcam). On day 7, representative hydrogels were also stained using a LIVE/DEAD assay to determine viability at the final time point. IHC specifically targeted insulin (1:200 dilution, ab181547, Abcam), LMN (1:200 dilution, ab11575, Abcam), and FN (1:200 dilution, ab2413, Abcam). Alexa Fluor secondary antibodies were used for fluorescent visualization.

### Bioink-based 3D adipose cultures

Adipocyte organoids were formed by placing preadipocytes (Lonza, PT-5020) in 10 μL of HA-Coll bioink at a cell density of 5 million cells/mL. The preadipocytes underwent a differentiation protocol in a 3D bioink culture in four different matrix compositions: a) HA-Coll, b) HA-Coll with FN, c) HA-Coll with LMN and d) HA-Coll with FN and LMN. FN and LMN were incorporated at 5 μg/mL. For adipose differentiation, preadipocyte-containing organoids were cultured in preadipocyte growth media (PromoCell, C-27410) for 4 days, then the preadipocyte growth media was switched to a preadipocyte differentiation media (PromoCell, C-27436) for 72 hours, and finally replaced with preadipocyte nutrition media (PromoCell, C-27438). The organoids were then maintained in culture for 12-14 days for full differentiation to occur, during which the nutrition media was replaced every 3 days. Extent of differentiation was visualized by staining the organoids with Nile Red (Invitrogen), which highlights lipids within the cells. Images were captured by macro-confocal microscopy (Leica TCS LSI, Leica Microsystems).

### 3D adipose functional testing

A high glucose insult study was performed to assess adiponectin response of adipocytes in HA-Coll+FN+LMN bioinks. The adipocyte media that the organoids are cultured in have a glucose content of 5 mM (Adipocyte nutrition media, PromoCell). This relatively low glucose media was removed from the adipocyte organoids, aliquoted, frozen, and stored before exposing them to high glucose (22 mM) for 48 hours. The high glucose media was then aliquoted, frozen, and stored, after which fresh 5 mM media was administered. After 48 hours the media was again aliquoted, frozen, and stored for further analysis. Collected media aliquots were thawed and analyzed quantitatively by ELISA for adiponectin (Abcam).

RNA was isolated from undifferentiated and differentiated fat spheroids using the RNeasy Fibrous Tissue Mini Kit (Qiagen, 74704). RNA was quantified with a Nanodrop 2000c (ThermoFisher Scientific). cDNA was synthesized using the High-Capacity cDNA Reverse Transcription Kit (ThermoFisher Scientific, 4368814). Reverse transcription reactions were processed as described by the manufacturer’s protocol at 25°C for 10 mins, 37°C for 2 hours and 85°C for 5 minutes. Quantitative PCR (qPCR) was performed using Applied Biosystems® QuantStudio^TM^ 3 Real-Time PCR System (ThermoFisher Scientific) with either SYBR® Green or TaqMan® probes. PCR primers and Taqman probe information is provided in ***Table S1***. All reactions were performed in replicates and normalized to *GAPDH* expression.

### Statistical analysis

The data are generally presented as the means of number of replicates ± the standard deviation. In most studies 3 to 6 replicates were employed. Unless otherwise described, data are graphed and analyzed for significance using a Student’s *t*-test or one way ANOVA. P-values below 0.05 were considered significant.

For the pancreatic islet insulin quantification study, an analysis considering the entirety of each curve was employed. Each group as a Gaussian curve was determined by a nonlinear regression analysis using GraphPad Prism. Subsequently, a one-way ANOVA was performed using the best-fit values of the regression analysis as means, standard deviation of the regression analysis as standard errors, and degrees of freedom of the regression analysis plus one as the number of replicates **n**. One-way ANOVA analysis was then performed, thereby considering the entire curves rather than individual time points.

## Acknowledgements

We are grateful for the administrative and project management support provided by Lynn Stedman, Brook Dohm, Pamela Shirling, Joshua Hunsberger, Kelly Burkett, Susan Lyons, and Julie Watson.

## Funding

The authors acknowledge funding from the Medical Technology Enterprise Consortium under Contract # W81XWH-15-9-0001. The views and conclusions contained herein are those of the authors and should not be interpreted as necessarily representing the official policies or endorsements, either expressed or implied of the U.S. Government. SH acknowledges funding from Research to Prevent Blindness. MW and AS acknowledge funding from the Wake Forest University and Wake Forest University Baptist Medical Center Cross Campus Pilot Program. AS also acknowledges funding from NIH grant R21CA229027, the Ohio State University, and the Ohio State University Comprehensive Cancer Center.

## Author contributions

AS conceived of the overall bioink platform and its components and properties. JA and AS oversaw and supervised all studies. AM, RCH, CC, and AM generated bioink formulations and performed rheology and printability testing studies. RCH, KY, and SB generated small molecule hydrogel modifiers. HS, TD, AM, RCH, KL, and YZ evaluated effectiveness of adhesion protein inclusion in bioinks. HS and SH performed liver bioink studies. HS and TD performed neural bioink studies. JA and YZ performed skeletal muscle studies. JA and AM performed pancreatic islet studies. HS and YZ performed adipose bioink studies. PG, MO, MW, TC, SS, and SH contributed to scientific discussion and evaluation of data. JA, HS, TD, YZ, AM, RCH, SH, and AS wrote the manuscript.

## Competing interests

The authors declare no competing interests.

## Data and materials availability

All data needed to evaluate the conclusions in the paper are included in the paper and/or the Supplementary Materials.

## Supplementary Materials

### Supplementary Materials Include

**Supplementary Figure 1.**
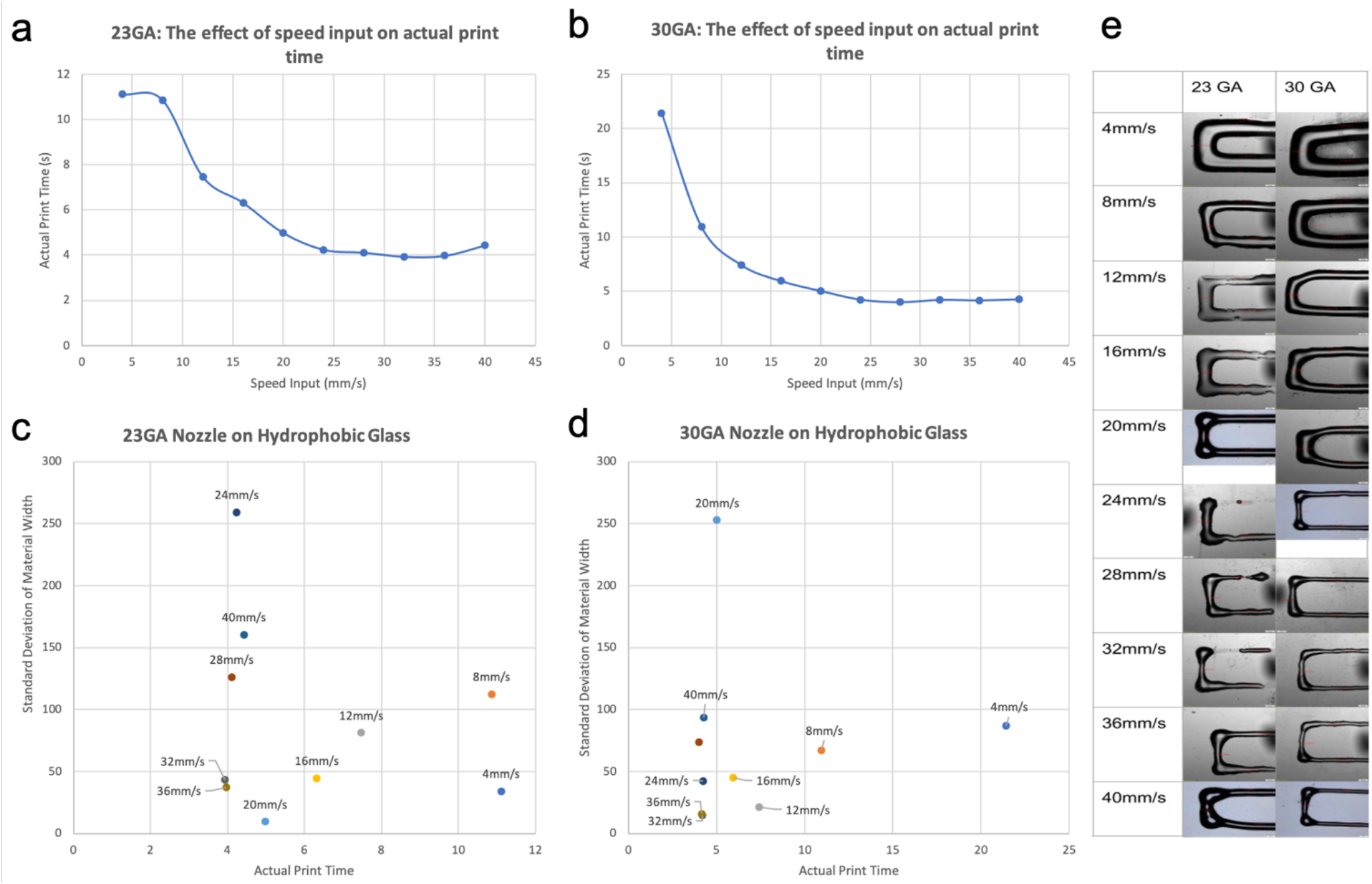
Printability assessment of the bioink using different gauge syringe tips and print speeds. a-b) Validation of nozzle-specific printing time by incremental speed increases. a) Increases in printing speed using a larger diameter nozzle, 23-gauge, decreases printing time at a stepping rate until reaching a plateau. b) A smaller nozzle, 30-gauge, has a more constant printing rate decrease as speed increases, with a similar plateau after 20 mm/s. c-d) At constant speeds, the width from the center of three perpendicular extruded lines were measured, the level of variance between each set indicates the consistency of line consistent line thickness. c) Larger 23-gauge nozzles maintain extrusion consistency at increased printing speeds and higher printing times. d) 30-gauge needles have a more constant resolution, lower standard deviation, at different printing speeds with no increased printing time. Ideal printing speeds for both nozzles between 32-36 mm/s. e) Representative images of the extruded perpendicular lines at different printing speeds with 23-gauge and 30-gauge nozzles.

**Supplementary Figure 2.**
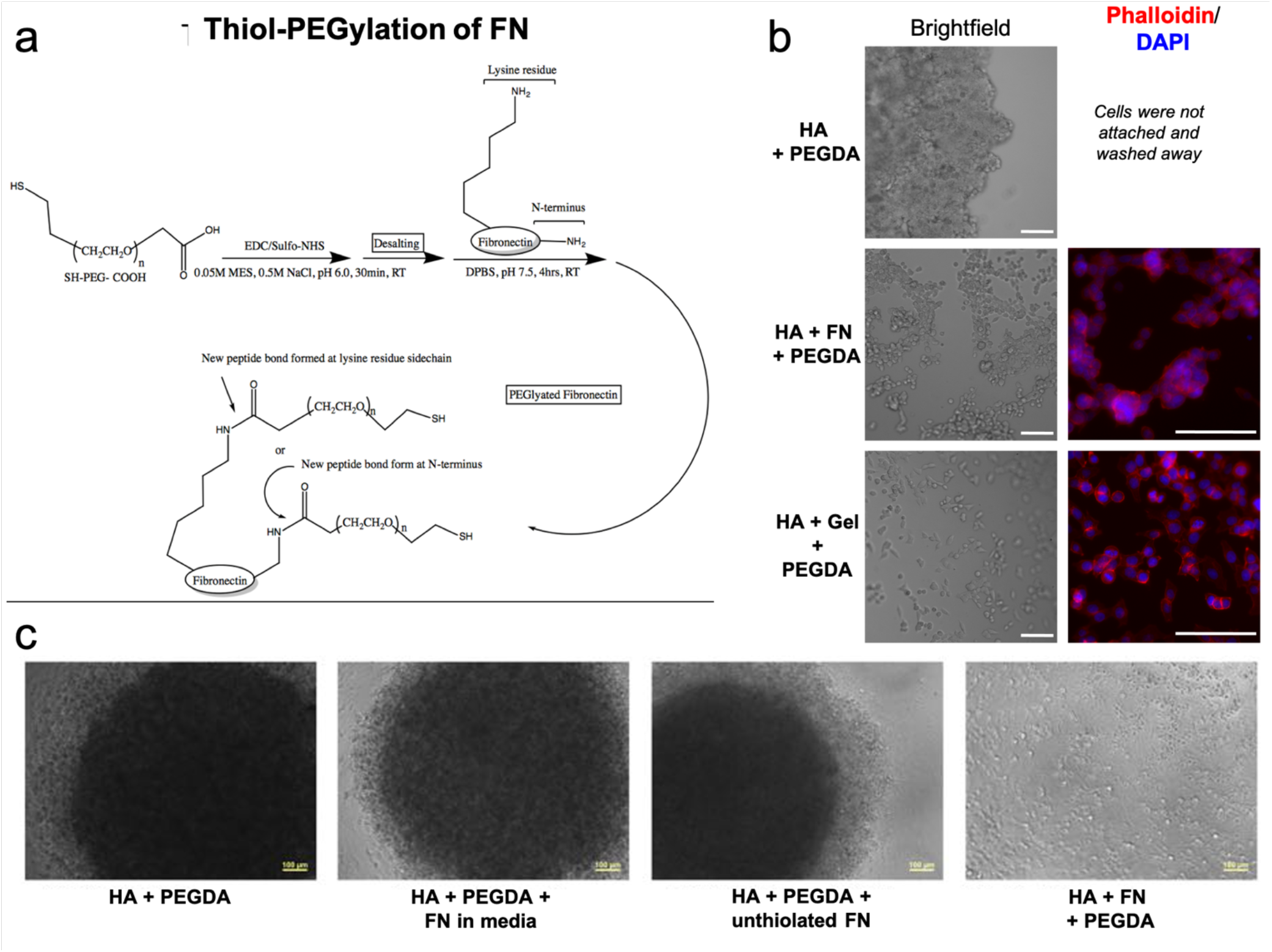
Detailed thiol-PEGylation of fibronectin and functional verification of successful fibronectin incorporation into HA hydrogels. a) Chemical synthesis scheme for modifying fibronectin with a thiol group for crosslinking into thiophilic hydrogels. b-c) Functional verification of covalent integration of modified fibronectin into hydrogels. b) Thiolated HA was prepared without methacrylated collagen to remove cell adhesion sites. Thiolated HA was crosslinked with PEGDA and supplemented with thiolated fibronectin or thiolated gelatin as control. Images show HCT-116 cells that aggregated on HA-PEGDA only hydrogels and then were washed away but adhered and spread out on hydrogels containing thiolated fibronectin or gelatin. c) HUVECs were placed in similar conditions, including a control hydrogel, a control hydrogel with fibronectin in media, and an unthiolated fibronectin incorporated in the hydrogel, both of which yielded no cell adherence to the hydrogel, but cell-cell aggregation. Only covalent incorporation of fibronectin into the hydrogel yielded adherence of cells at the surface of the hydrogel substrate.

**Supplementary Figure 3.**
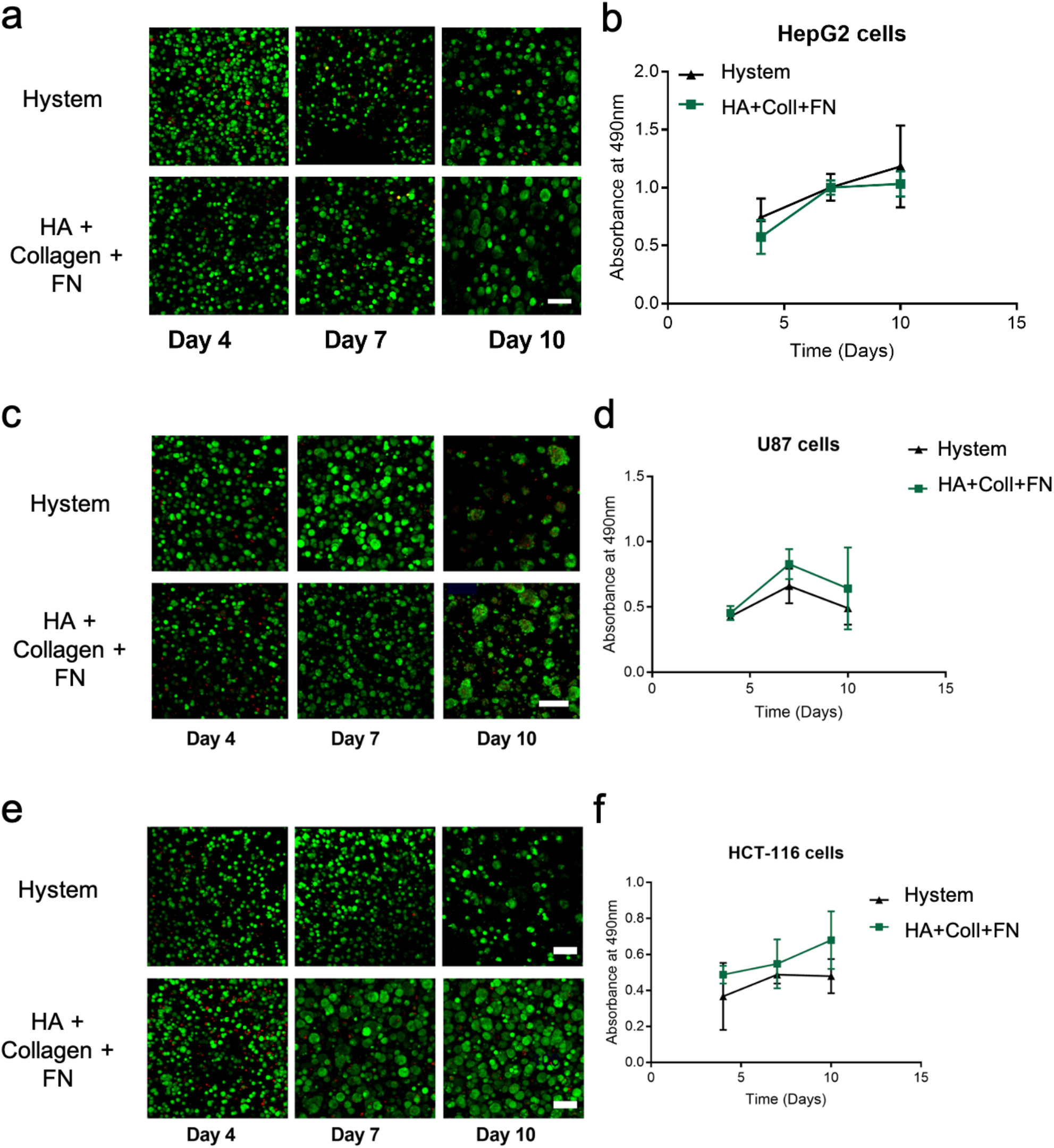
Cell line verification that synthetically modified adhesion protein (fibronectin) does not cause detrimental effects when integrated into the HA-Coll bioink. LIVE/DEAD viability/cytotoxicity assays and proliferation by ATP activity quantification were performed for commonly used cell lines-b) HepG2 hepatoma, c-d) U87 MG glioma, and e-f) HCT-116 colorectal cancer cells cultured within an established control hydrogel (commercially available Hystem, a thiolated HA, thiolated gelatin, and PEGDA hydrogel) or the HA-Coll bioink supplemented with thiolated fibronectin. a, c, d) Green – Calcein AM-stained viable cells; Red – Ethidium homodimer-1-stained dead cell nuclei. Scale bars – 100 μm. b, d, f) No significant differences were observed between groups.

**Supplementary Figure 4.**
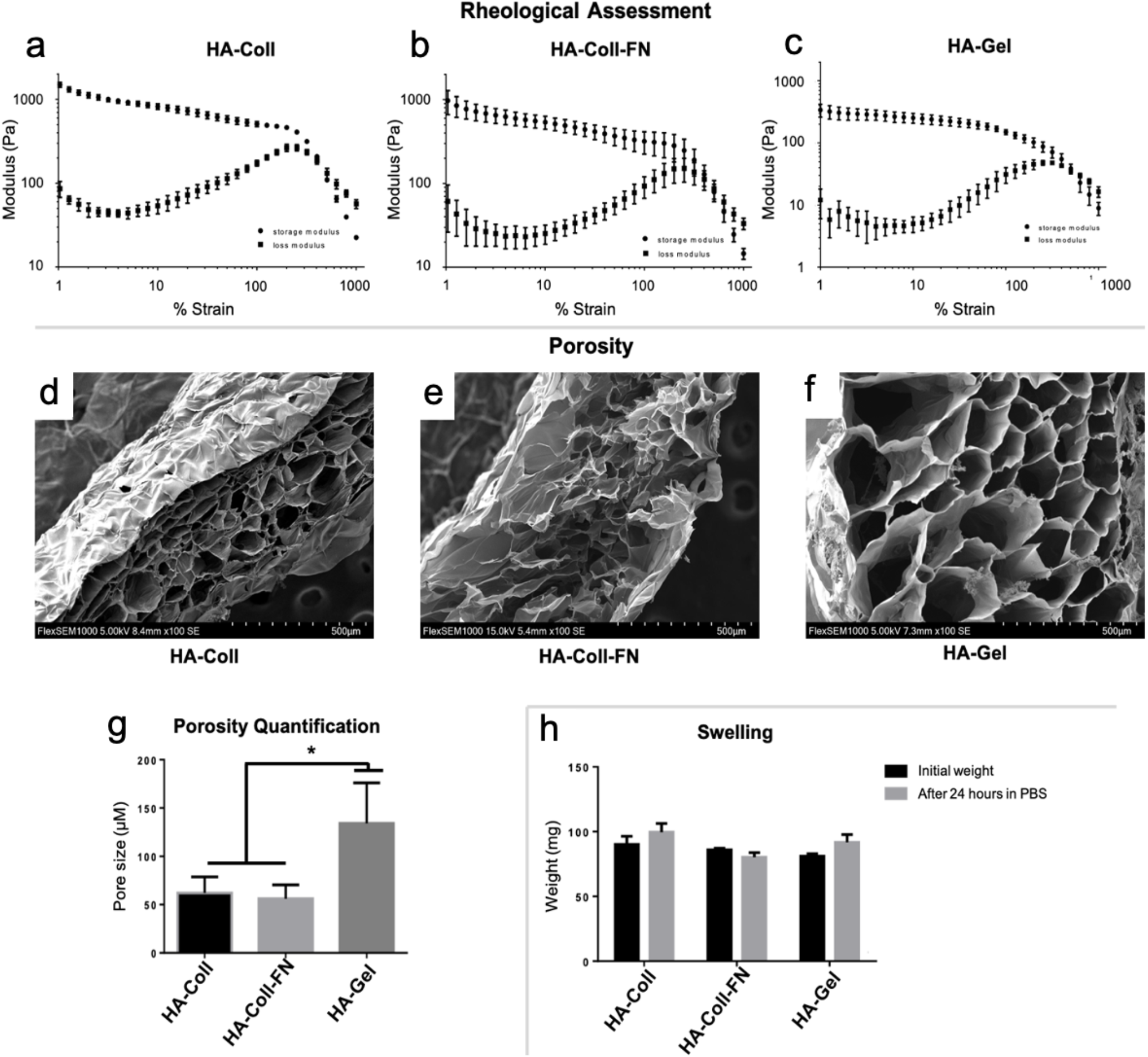
Hydrogel characterization. a-c) Rheological assessment of hydrogels. Shear elastic (storage) modulus, G’, and shear loss modulus, G”, are shown in response to strain sweeps (0 – 1000%) for a) HA-Coll, b) HA-Coll-FN, and c) HA-Gelatin hydrogels. d-f) SEM images show internal porosity of d) HA-Coll, e) HA-Coll-FN, and f) HA-Gelatin hydrogels. g) Quantification of pore diameter. Statistical significance: * p < 0.05. h) Swelling characteristics of the hydrogels after incubation with PBS for 24 hours.

**Supplementary Figure 5.**
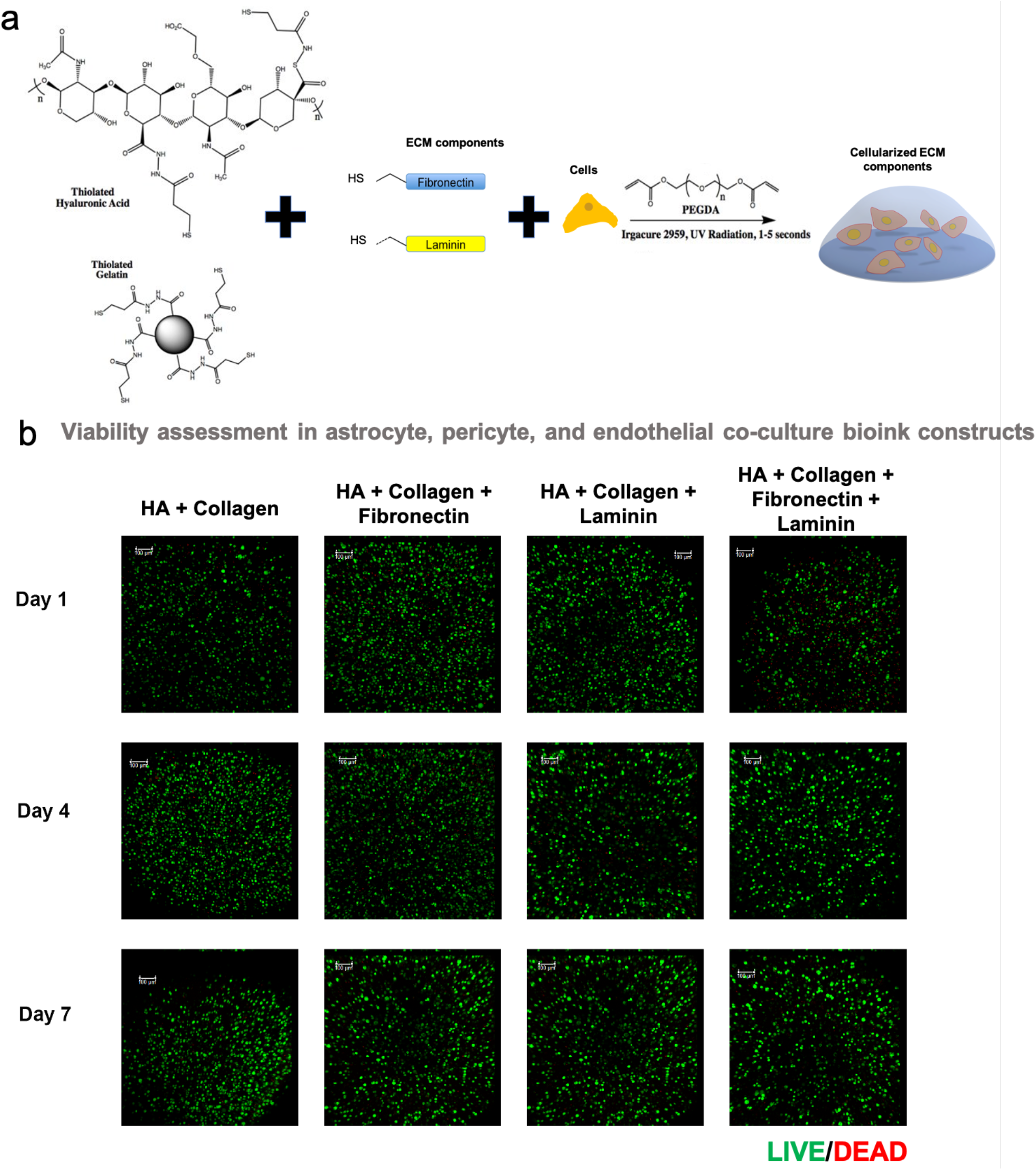
Initial evaluation of hydrogel bioink modulation on heterogeneous neural constructs. a) Schematic of hydrogel modulation, formulation, and crosslinking. Thiolated hyaluronic acid, thiolated gelatin, thiolated adhesion proteins, PEGDA, and cells (astrocytes, pericytes, and endothelial cells) are mixed, after which in the presence of a photoinitiator brief UV light irradiation induces crosslinking. B) Viability assessment of neural constructs by LIVE/DEAD staining and macro-confocal imaging. Green – Calcein AM-stained viable cells; Red – Ethidium homodimer-1-stained dead cell nuclei. Scale bars – 100 μm.

**Supplementary Figure 6.**
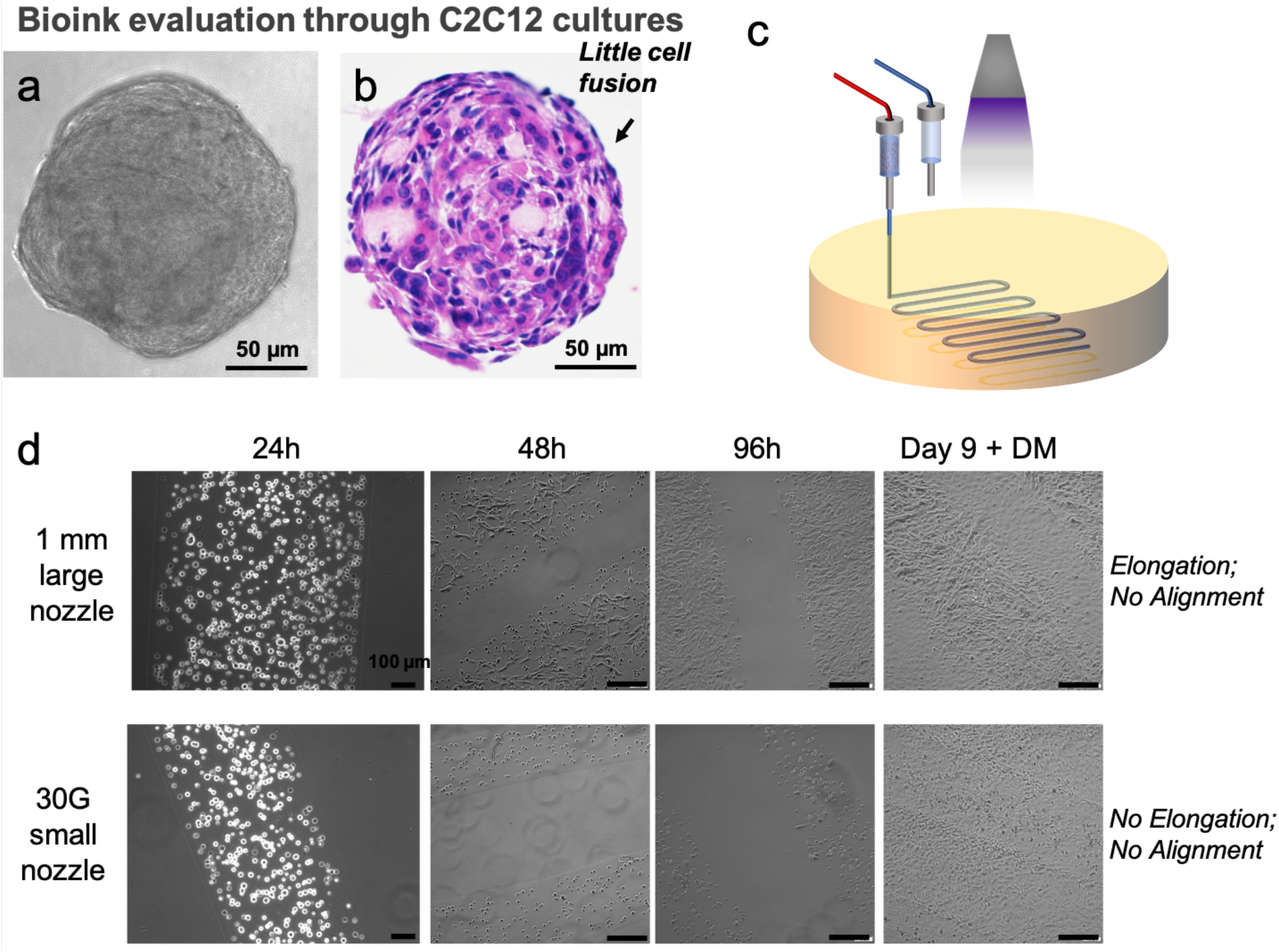
Evaluation of using bioprinting bioink filaments to align C2C12 cells. a-b) C2C12 spheroid cultures, visualized by a) light microscopy and b) hematoxylin and eosin staining, show little evidence of cell fusion. c-d) Exploration of using bioprinted bioink filaments to drive cellular alignment and fusion. c) Schematic of filament bioprinting, using UV crosslinking to stabilize bioprinted bioink conformations. D) Light microscopy images of C2C12 cells within bioprinted bioink filaments over time, and after introduction of differentiation media DM). Scale bars – 100 μm.

**Supplementary Figure 7.**
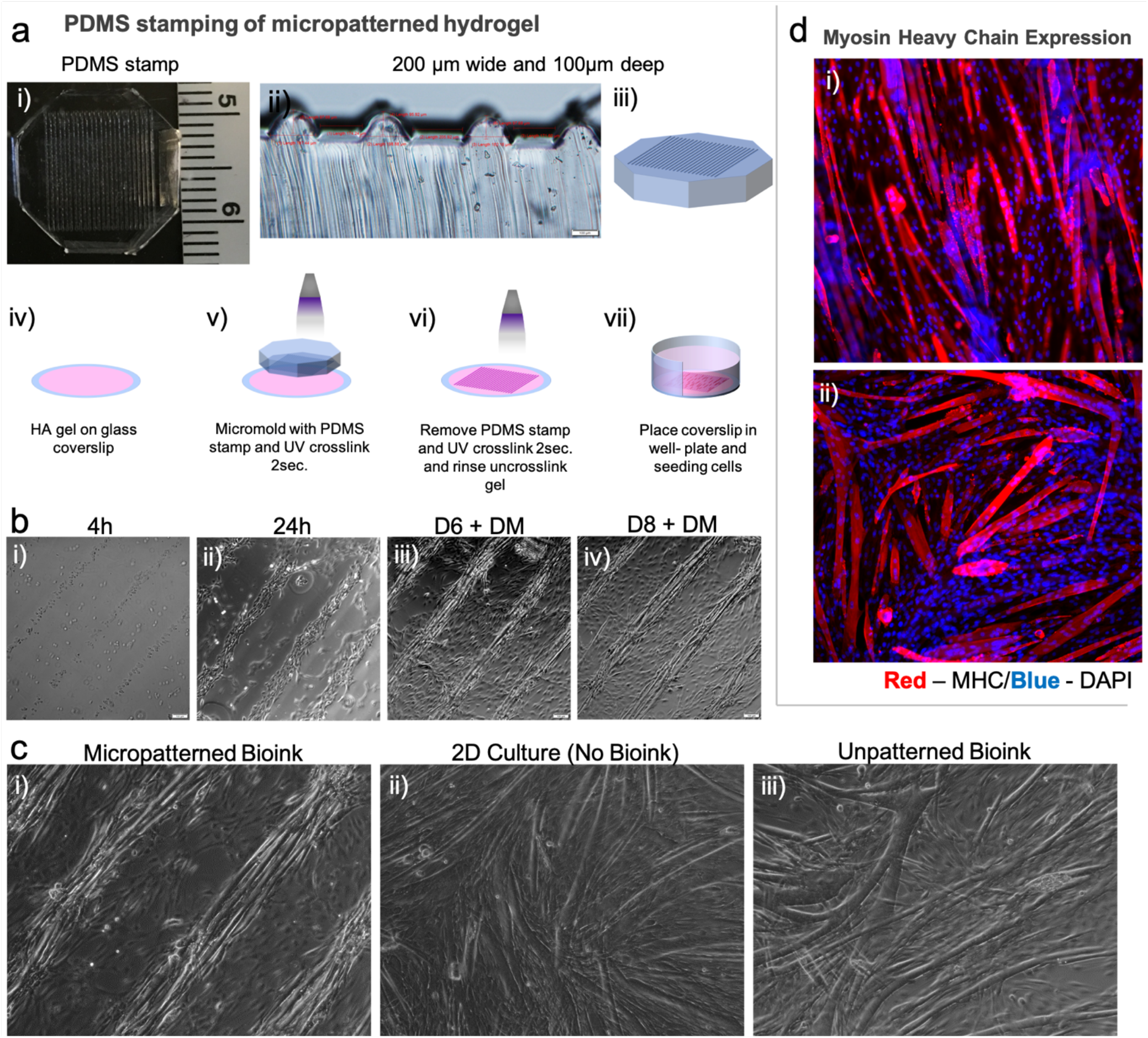
Evaluation of topographical patterning of HA bioinks to induce alignment and fusion of C2C12 cells. a) Topographical stamping was enabled by micropattern molding of a i) polydimethylsiloxane (PDMS) stamp with ii) microscale grooves. iii) The PDMS stamp was then placed on the iv) uncrosslinked HA bioink surface, v) UV crosslinked, vi) the stamp was removed and the bioink was further crosslinked and rinsed, and vii) finally the micropatterned bioink substrate was transferred to well-plates for cell studies. b) C2C21 cells align over time and fuse when cultured with skeletal muscle differentiation media (DM). c) Comparison of alignment and fusion between differentiated C2C12 cells on i) micropatterned bioinks, ii) in standard 2D plastic culture, and iii) on unpatterned bioinks. d) Immunofluorescent staining of myosin heavy chain in C2C12 cells on i) micropatterned bioinks versus ii) unpatterned bioinks. Red – myosin heavy chain; Blue – DAPI.

**Supplementary Figure 8.**
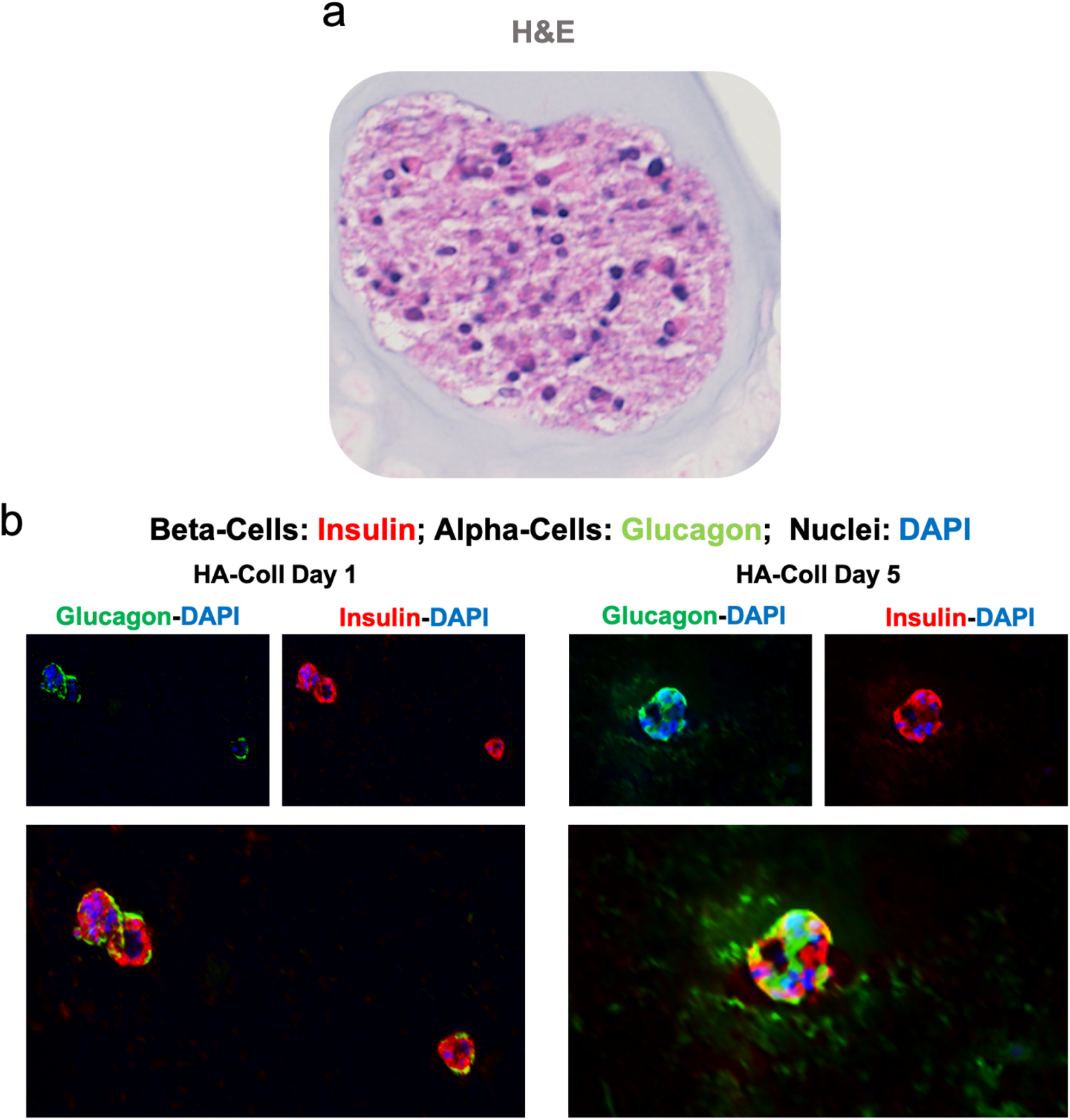
Characterization of HA-Coll bioink-supported human pancreatic islets. a) Hematoxylin and eosin staining of human pancreatic islets bioprinted in HA-Coll bioinks. b) Immunofluorescent staining in 3D of HA-Coll bioprinted islets on day 1 and day 5 following bioprinting. Green – glucagon; Red – insulin; Blue – DAPI.

**Supplementary Figure 9.**
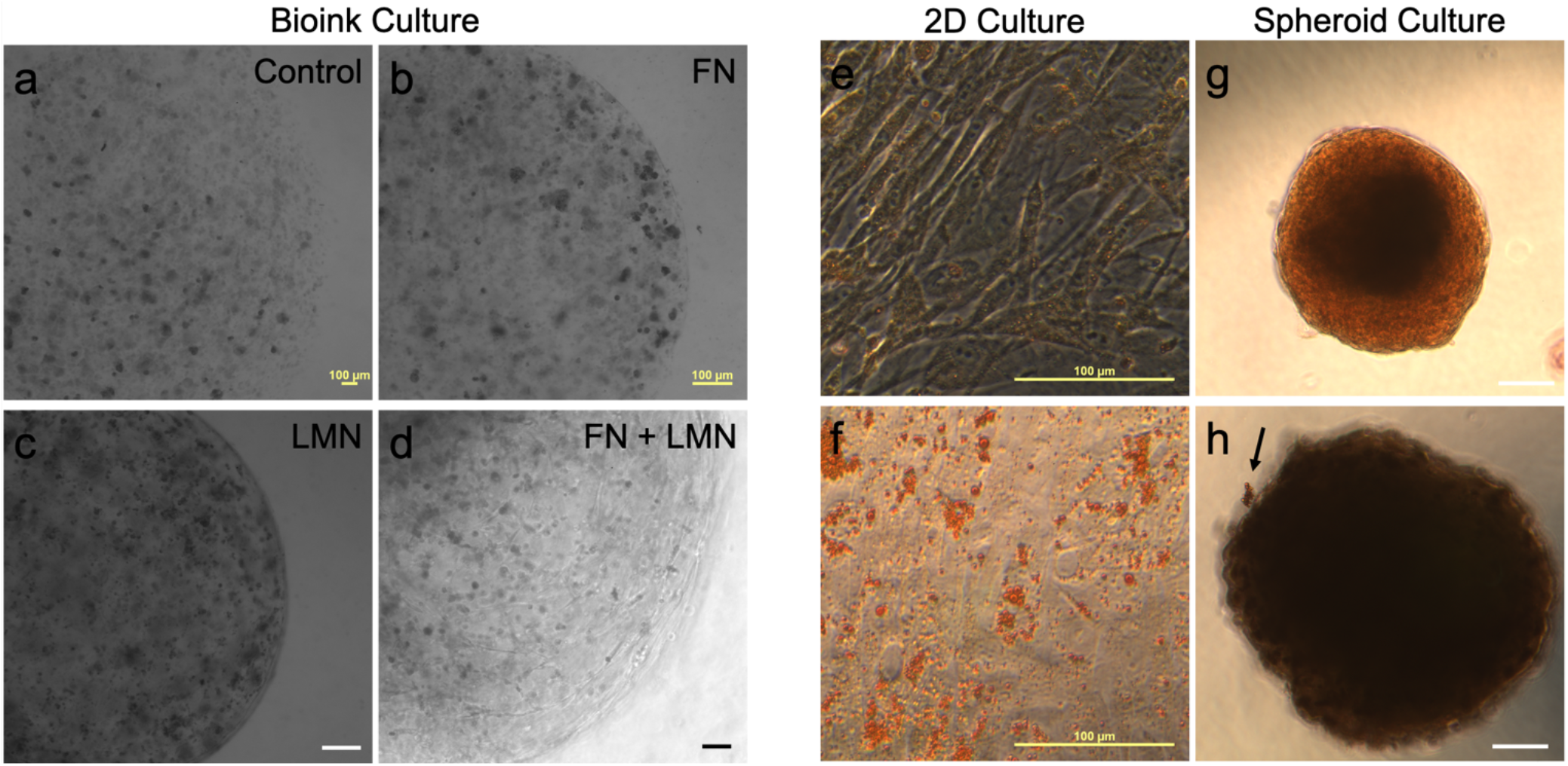
Human preadipocyte 3D construct cultures in HA-Coll bioinks. a-d) HA-Coll (control), HA-Coll+FN, HA-Coll+LMN, and HA-Coll+FN+LMN constructs visualized by light microscopy. e-f) Assessment of Nile Red staining in 2D versus 3D structures. Scale bars – 100 μM.

**Supplementary Figure 10.**
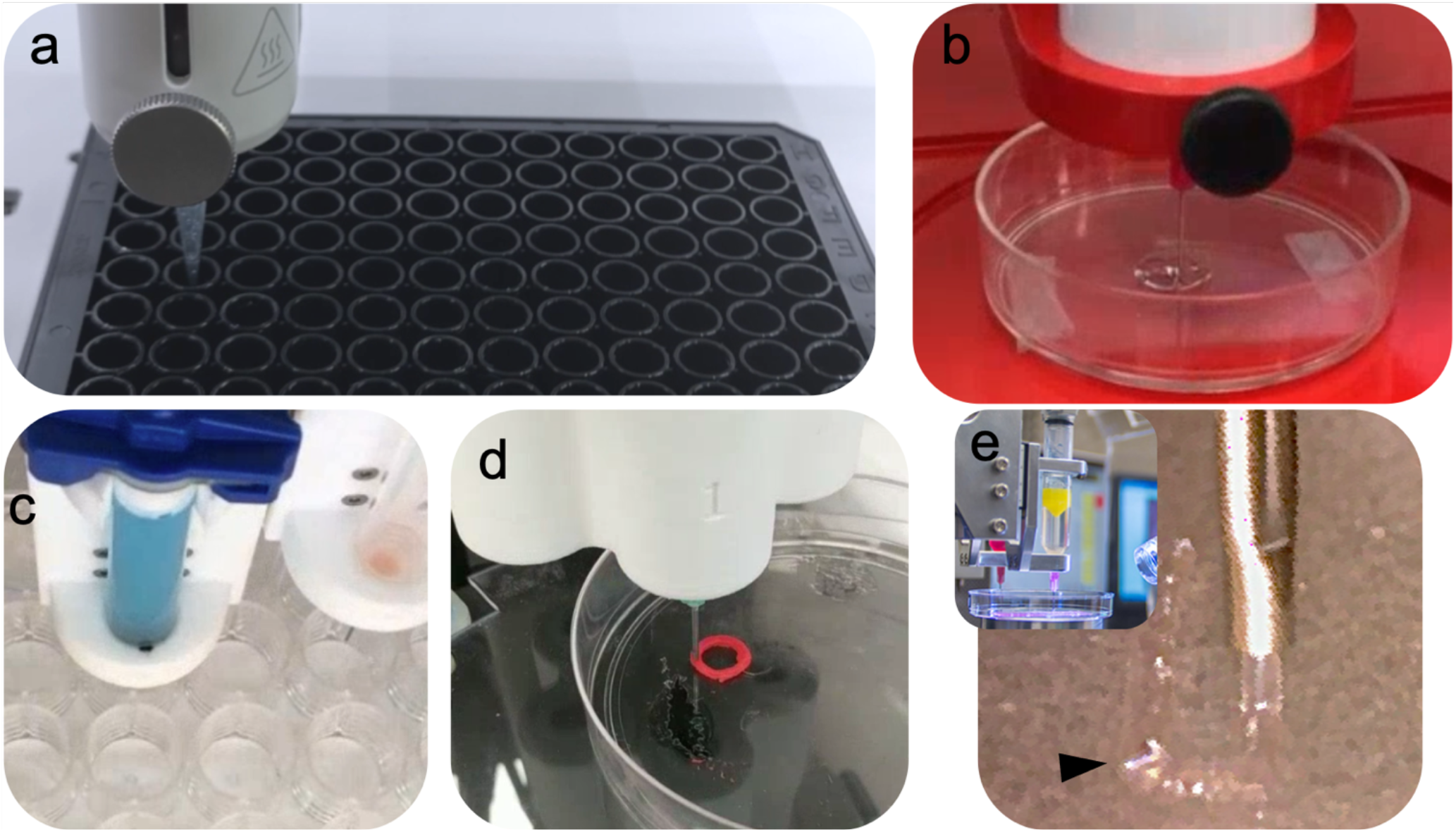
HA-Coll bioink bioprinting on board a variety of bioprinter platforms. We have successfully deployed our bioink technology on the a) CELLINK BIO X, b) Allevi Allevi2, c) CELLINK Inkredible, d) Advanced Solutions Biobot, and e) the Wake Forest ITOP (Integrated Tissue-Organ Printer).

**Supplementary Table 1:**
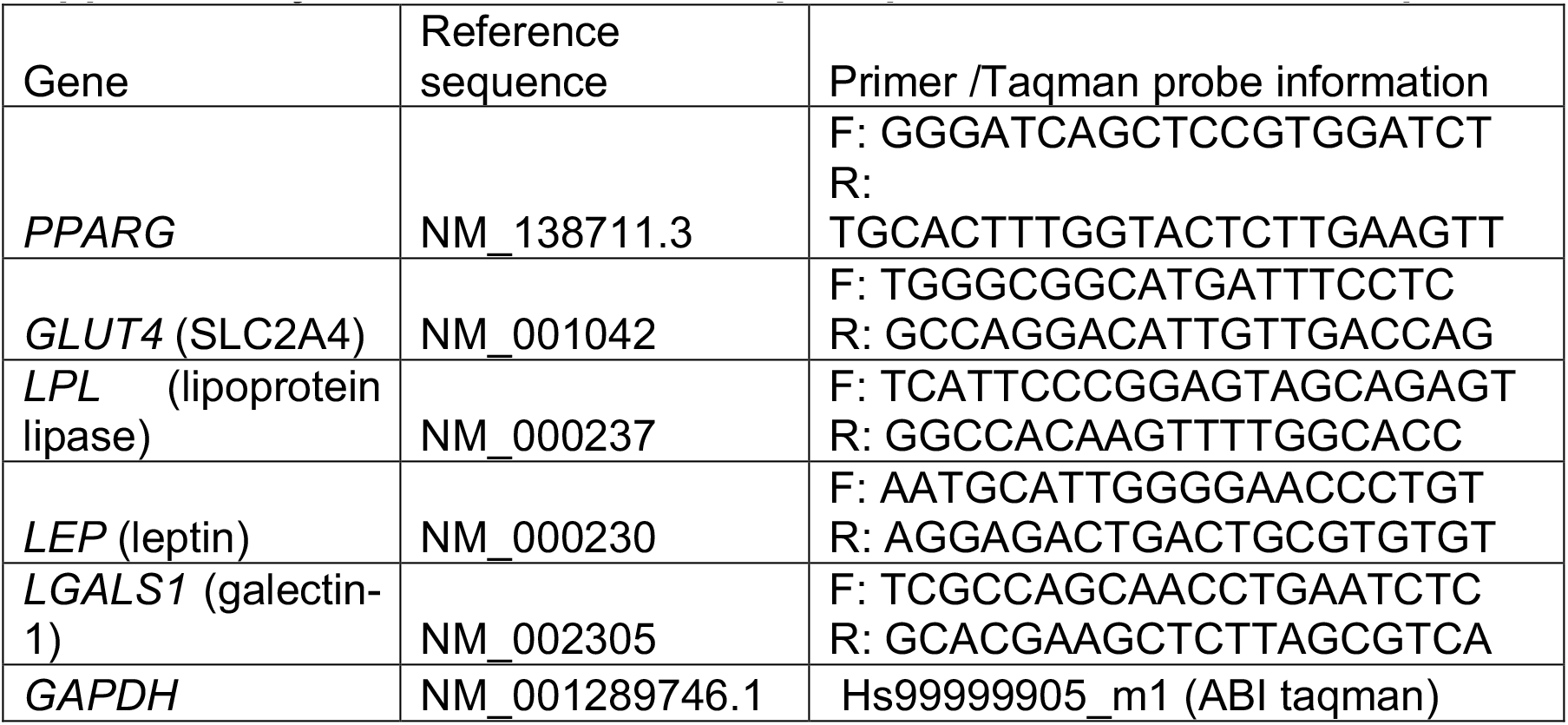
Primer or TaqMan probe information for adipose function test.

## Notes

### Competing Interest Statement

The authors have declared no competing interest.

